# Shoot at Site: Advancing *in planta* transformation, regeneration and gene-editing through a cascade of wounding-mediated developmental regulators

**DOI:** 10.1101/2025.02.06.636944

**Authors:** Arjun Ojha Kshetry, Kaushik Ghose, Anshu Alok, Vikas Devkar, Vidhyavathi Raman, Robert M. Stupar, Luis Herrera-Estrella, Feng Zhang, Gunvant B. Patil

## Abstract

Developing transgenic and/or gene-edited plants largely depends on tedious, lengthy, and costly *in vitro* regeneration protocols. While plants have remarkable regeneration ability, not all species, genotypes or even explants exhibit the same transformation and regeneration potential under *in vitro* conditions. To tackle this bottleneck, we have developed a seamless and user-friendly system to induce transgenic and gene-edited *de novo* meristems via a synthetic cascade comprising a wound-induced regeneration pathway, plant developmental regulators (DRs) and gene-editing reagents. *WOUND INDUCED DEDIFFERENTIATION 1 (WIND1)* is used as a transcriptional regulator to control the expression of various DR genes through *ENHANCER OF SHOOT REGENERATION 1 (ESR1)* promoter. This cascade was strategically applied *in planta* to the non-meristematic internode of *N. benthamiana* to induce meristematic activity and regenerate *de novo* shoots with knock-out mutations of the *phytoene desaturase* (*PDS*) gene. This synthetic toolkit was further applied successfully to tomato and soybean. This methodology offers a transformative approach to overcome barriers in plant biotechnology, potentially accelerating the generation of transgenic and gene-edited plants without reliance on conventional tissue-culture intermediates.

## Main

Plant tissue culture or *in vitro* regeneration is a cornerstone in genetic engineering and plant biotechnology. However, only a limited number of plant species exhibit amenable characteristics for this process, often requiring several months to achieve successful regeneration [1]. Additionally, the hurdle in obtaining transgenic and gene-edited progeny not only relies on their regeneration plasticity but also the successful transformation of the transgene and gene-editing reagents which significantly impede the efficient development of transgenic and gene-edited plants [2, 3]. Somatic cell regeneration is a highly complex process and success in transformation and regeneration is influenced by several pivotal factors, encompassing the application of plant growth regulators, the formulation of the basal media, the nature of the explant, selectable markers and transformation methods [4, 5]. Significantly, this process depends on the species and genotype, underscoring the importance of precision in understanding and manipulating these variables. Considerable efforts have been dedicated to investigating innovative approaches for regeneration and transformation, aiming to overcome the conventional procedures dependent on tissue culture-based transformation [6, 7]. Early endeavors to adopt a “tissue culture-free” transformation approach involved the use of *Agrobacterium* to transform germinating seeds [8] and *in planta* transformation and regeneration of apical shoots in Arabidopsis [9]. Later, a simple floral dip transformation method was developed in Arabidopsis to successfully generate transgenic events [10]. However, this method is suitable to only a few species, mainly belonging to the *Brassicaceae* family. In parallel to these endeavors, several researchers have identified that the ectopic expression of morphogenic genes or developmental regulators (DRs) such as *isopentenyl pyrophosphate transferase* (*ipt*), *WUSCHEL* (*WUS*), *BABY-BOOM* (*BBM*), *SHOOT MERISTEMLESS* (*STM*), among others, dramatically enhanced the efficiency of plant transformation and regeneration [11–15]. More recently, with the advent of gene-editing technology, the concept of tissue culture-free transformation in plants became even more critical for the general application of this technology in agriculture. Tissue culture-free plant gene editing has been achieved by the co-delivery of gene editing reagents (CRISPR/Cas9 and guide RNAs) with DRs that induce *de novo* organogenesis, thereby obtaining stable targeted gene-edited plants [6, 16]. However, oftentimes expression of DRs alone is not enough for direct organogenesis.

They must collaborate with endogenous or exogenous phytohormones, wound stress, plant developmental stages and several other factors, thereby establishing and maintaining meristematic activity and reprogramming of differentiated somatic cells [6, 14, 17, 18].

Among several regeneration pathways, developing *de novo* organs induced by wounding stress is one of the primary triggers of plant regeneration (**Supplementary Fig. 1**). This property of wounding and wound healing has long been utilized for clonal propagation such as cutting and grafting [19, 20], and direct and indirect somatic cell regeneration [21]. In nature, plants are constantly challenged by harsh environments, predators, diseases, and physical injury, but they overcome these diverse challenges by repairing local wounds and reconstructing tissues or organs [22]. The wound-healing process induces the transcriptional activation of molecular signals necessary for reprogramming, allowing local cells to heal, survive, and initiate new growth and development [23, 24]. Additionally, it triggers the biosynthesis of endogenous hormonal pathways, crucial for determining cellular fate during regeneration [5, 25]. Although, the development of masses of undifferentiated cells or callus after wounding is the first step of tissue healing, appropriate biosynthesis of endogenous hormones and activation of molecular signals can induce cell fate reprogramming and initiate meristematic activity [21, 26]. Notably, during this process, a wound-induced tissue repair pathway gets activated involving transcriptional regulators belonging to *APETAL2/ETHYLENE RESPONSE FACTOR* (*AP2/ERF*) gene family; *WOUND INDUCED DEDIFFERENTIATION 1 (WIND1), ENHANCER OF SHOOT REGENERATION 1 (ESR1)*, and *PLETHORA3 (PLT3*), *TYPE-B ARABIDOPSIS RESPONSE REGULATORS* (*ARRs*), which reprogram cell fate and initiation of new developmental pathways [21]. Among these regulators, *WIND1* is a master regulator of wound-induced cellular reprogramming in plants. It promotes callus formation, formation of tracheary elements, and pluripotency acquisition as a primary step for plant regeneration [27–30].

Following wounding, the expression of *WIND1* is upregulated within 30 minutes and peaks to about ten-fold one hour after wounding, which in turn acts as a transcriptional activator of *ESR1* belonging to the *AP2/ERF* gene family [27]. Furthermore, ectopic expression of *ESR1* in Arabidopsis induces callus formation and initiates shoot regeneration under optimal *in vitro* conditions [27, 31, 32].

Previously, several studies have employed various hormone biosynthesis genes and DRs to enhance the efficiency of genetic transformation and plant regeneration [6, 11–14]. Lowe et al., (2016), significantly improved transformation efficiency in recalcitrant maize inbreds through expressing *BBM* and *WUS2* genes [14]. Similarly, Debernardi et al., (2020) demonstrated enhanced *in vitro* transformation of rice, wheat, and citrus by expressing a *GRF-GIF* chimeric protein [18]. However, these approaches still depend on long, laborious, and complex tissue-culture protocols. In recent advancements, Maher et al. (2020) and Lian et al. (2022) developed tissue-culture-free methods for plant regeneration and gene editing by overexpressing combinations of DR genes in various plant species. Similarly, viral delivery methods have been employed for heritable, tissue-culture-free gene editing in plants like tomatoes and tobacco [16, 33]. However, these approaches often require plants to mature, involve DR delivery at axillary meristems, and demand extended periods to achieve *de novo* shoot formation. Additionally, regenerated plants frequently exhibit severe pleiotropic effects or depend on viral vectors specific to plant species, highlighting the need for improved strategies [34]. We developed an innovative approach leveraging the *WIND1-ESR1* cascade to address these challenges and accelerate *de novo* shoot regeneration. We successfully developed transgenic and gene-edited plants by transfecting young tobacco and tomato plants (under one month old) at internodes, and embryo axes of soybeans lacking meristematic activity. This method enabled shoot induction as early as two weeks post-transfection, significantly enhancing regeneration efficiency. In this study, we utilized the wound-response pathway to overcome limitations associated with labor-intensive tissue culture, prolonged regeneration times, and pleiotropic effects (**Supplementary Fig. 1**). We engineered robust synthetic cascade coordinating genes involved in stem cell maintenance, rapid tissue differentiation, and regeneration under wound-induced regulatory control. Important DR genes, including *ipt*, *WUS, STM, BBM,* and the *GRF-GIF* chimera, were individually tested to assess *de novo* shoot formation in tobacco, tomato, and soybean.

Additionally, this system was combined with gene-editing reagents to target the *phytoene desaturase* (*PDS*) gene in tobacco and soybeans. This streamlined approach offers a promising framework for advancing efficient, scalable, and cost-effective plant regeneration and gene editing.

## RESULTS

### Assessment of *WIND1:ESR1* response

The *WIND1* plays a central role in promoting wound-induced cellular reprogramming in plants, a process crucial for callus formation through the activation of cytokinin response [27]. One of the critical outcomes of *WIND1* activity is transcriptional activation of downstream regulator genes, such as *ESR1,* guiding somatic cells toward a dedifferentiated state (wound healing) and in some cases, initiation of organogenesis {**Supplementary Fig. 1**; [27, 32]}. With this reasoning and to test the cascade effect of *AtWIND1* on the promoter of *AtESR1* (pro*AtESR1*) in tobacco (*N. benthamiana*), we designed a series of cascade constructs (**Fig. 1** and **Supplementary Table S1)**. The *RUBY* gene [35] was used to visualize transgene expression non-invasively. The ectopic expression of the *RUBY* gene results in betalain accumulation, causing bright red pigmentation in plant tissues. The construct proCmYLCV::*RUBY* and pro*AtESR1::RUBY* were used as controls to assess callus formation and gene induction under the control of *AtESR1*, respectively. Finally, to verify if *AtWIND1* binds and activates the *AtESR1* promoter and induces callus formation, we developed a construct carrying pro35S::*WIND1*-pro*ESR1*::*RUBY* cassettes. As expected, relatively low callus formation was observed when the *RUBY* gene was expressed under proCmYLCV or the *AtESR1* promoter in tobacco explants, due to the natural response of wounding during explant preparation (**Fig. 1A and B**). However, explants transformed with pro35S::*AtWIND1* demonstrated a 3-4-fold increase in callus formation in phytohormone-free MS media (**Fig. 1C**).

**Figure 1:**
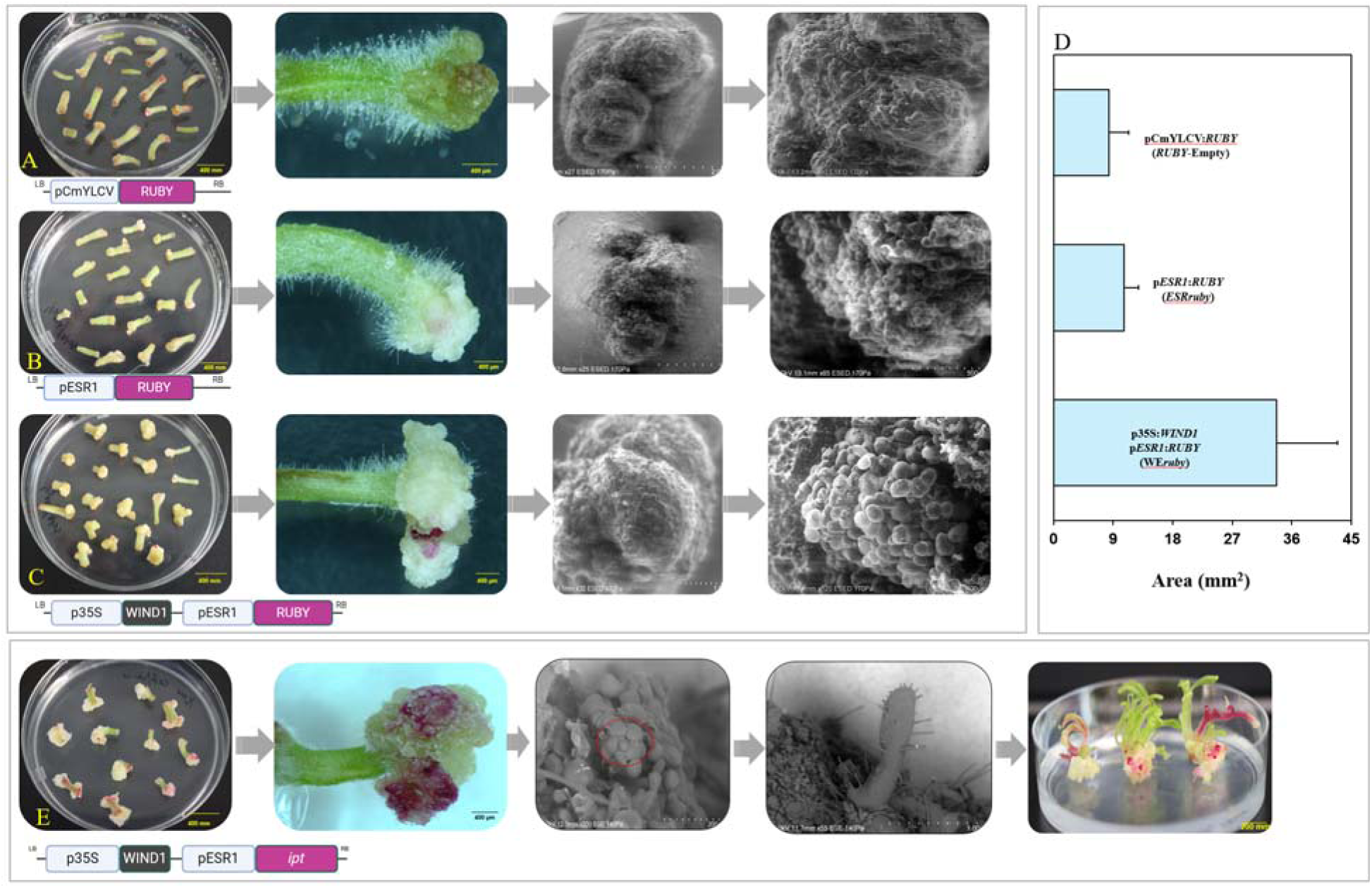
Experimental validation of gene constructs designed to assess p*ESR1*-driven gene activation in response to *AtWIND1* expression. Tobacco shoot segments transformed with *Agrobacterium tumefaciens* (GV3101) containing CmYLCV::*RUBY* **(A)** and p*At*ESR1::*RUBY* **(B)** showed limited or no callus formation. Whereas co-expression of 35S::*AtWIND1 – AtESR1::RUBY* **(C)** showed significantly more callus formation and activation of the *AtESR1* promoter. **(D)** Quantitative analysis of callus formation, shown in Area (mm^2^). Explants transformed with CaMV 35S::*AtWIND1* showed approximately 3-4-fold more callus formation. **(E)** Expression of developmental regulator genes like *ipt* under the *AtESR1* promoter and its activation by *AtWIND1* induced rapid callus induction, and formation of shoot apical meristems, leading to the induction of *de novo* meristems in phytohormone-free media.

These results validated that the overexpression of *AtWIND1* in tobacco accelerates the callus formation, activating the gene underlying the *AtESR1* promoter. Therefore, we reasoned that driving the expression of DR genes under the *AtESR1* promoter could enhance *de novo* shoot formation at the wound site. To test this hypothesis, we designed and constructed several vectors. In these constructs, DR genes were driven by the *AtESR1* promoter, and *AtWIND1* was driven by the CaMV 35S promoter, alongside *RUBY* driven by CmYLCV promoter as a visible reporter gene. Given *ipt* has been identified as a key cytokinin biosynthesis gene capable of promoting *de novo* shoot formation [6, 11], we first tested the *WEipt* construct under tissue culture conditions using internode and petiole explants of tobacco. Within three weeks, the *WEipt* significantly enhanced callus formation and induced the formation of multiple *de novo* shoot meristems on phytohormone-free ½ strength MS media (**Fig. 1E**). This successful *in vitro* regeneration confirmed that controlled expression of *ipt* under pro*AtESR1* mediated by *AtWIND1*, can effectively induce *de novo* shoot regeneration (**Fig. 1E**). This outcome provided a robust foundation for further testing various DR genes for *in planta* transformation experiments.

### *In planta* shoot formation, transgenesis and gene editing

To assess the combined effect of *AtWIND1* and pro*AtESR1* with different DRs on *de novo* meristem formation and *in planta* transgenesis and gene-editing, several well characterized DRs including *AtWUS, AtSTM, ipt, ZmBBM,* and *GRF-GIF* chimera were cloned under pro*AtESR1* in one of the modular vectors. Additionally, modular vectors containing *AtWIND1*, *RUBY*, and gRNAs targeting the tobacco *Phytoene Desaturase* (*PDS*) gene were constructed (**Supplementary Table 1**). These constructs were then introduced into young, Cas9-expressing transgenic tobacco plants by targeting three simple delivery sites: (1) multiple internode sites, (2) internodal areas proximal to the shoot apical meristem, and (3) pruned shoot sites (**Fig. 2B and Supplementary Fig. 2**). Following the *Agrobacterium* delivery, injection sites were covered with cotton plugs (soaked in the same *Agrobacterium* solution prepared for transfection), creating a localized microenvironment conducive to co-cultivation. These plants were incubated in the dark at about 80% humidity and 22°C for three days. The injection sites were carefully observed for callus formation and initiation of *de novo* buds. Since the *RUBY* gene was incorporated with all DRs genes, the expression of *RUBY* (red pigment) and the development of callus and *de novo* shoot formation were recorded after 1 week of transformation. Notably, all *AtWIND1*-expressing constructs, regardless of the delivery site, accelerated callus formation at the injection site compared to constructs without *AtWIND1*. The *de novo* shoot formation did not occur when constructs were delivered at multiple internode sites (**delivery site 1**) or areas proximal to the apical shoot (**delivery site 2**). For instance, injections at multiple sites along each internode led to callus formation, but the calli at these sites were more compact and exhibited signs of wound healing (**Supplementary Fig. 2**). Similarly, injections proximal to the shoot apical meristem resulted in compact callus formation, albeit relatively larger than multiple injection sites. In contrast, constructs carrying *AtWIND1* and *ipt* genes injected into pruned young shoots (**delivery site 3**) of *N. benthamiana* led to not only callus formation but dramatically accelerated the pace and frequency of *de novo* shoot formation at the wound site, as early as 12 days after transformation (**Fig. 3 and Supplementary Fig. 2).** The regeneration frequency using *ipt* was ∼71%. Expression of *GRF-GIF* on pruned shoots resulted in induction of *de novo* meristems at the injection sites, however, the efficiency was less than 4%, and no transgenic shoots were obtained (**Fig. 3A**). DR constructs with *AtWUS*, *AtSTM*, and *ZmBBM* did not result in *de novo* shoot formation, and their response was like the empty vector. Following the successful regeneration of *de novo* shoots at site of injection in tobacco with the third method, we employed similar delivery sites and *in planta* transformation strategy for tomato and soybean (**Fig. 2C and D**). Previous *in planta* transformation methods used mature plants (around 65-70 days old), and it also involved repeated axillary shoot removal, or multiple *Agrobacterium* injections [6, 13]. Our approach used younger, healthy seedlings (22–28 days old) to accelerate *de novo* shoot regeneration efficiency.

**Figure 2:**
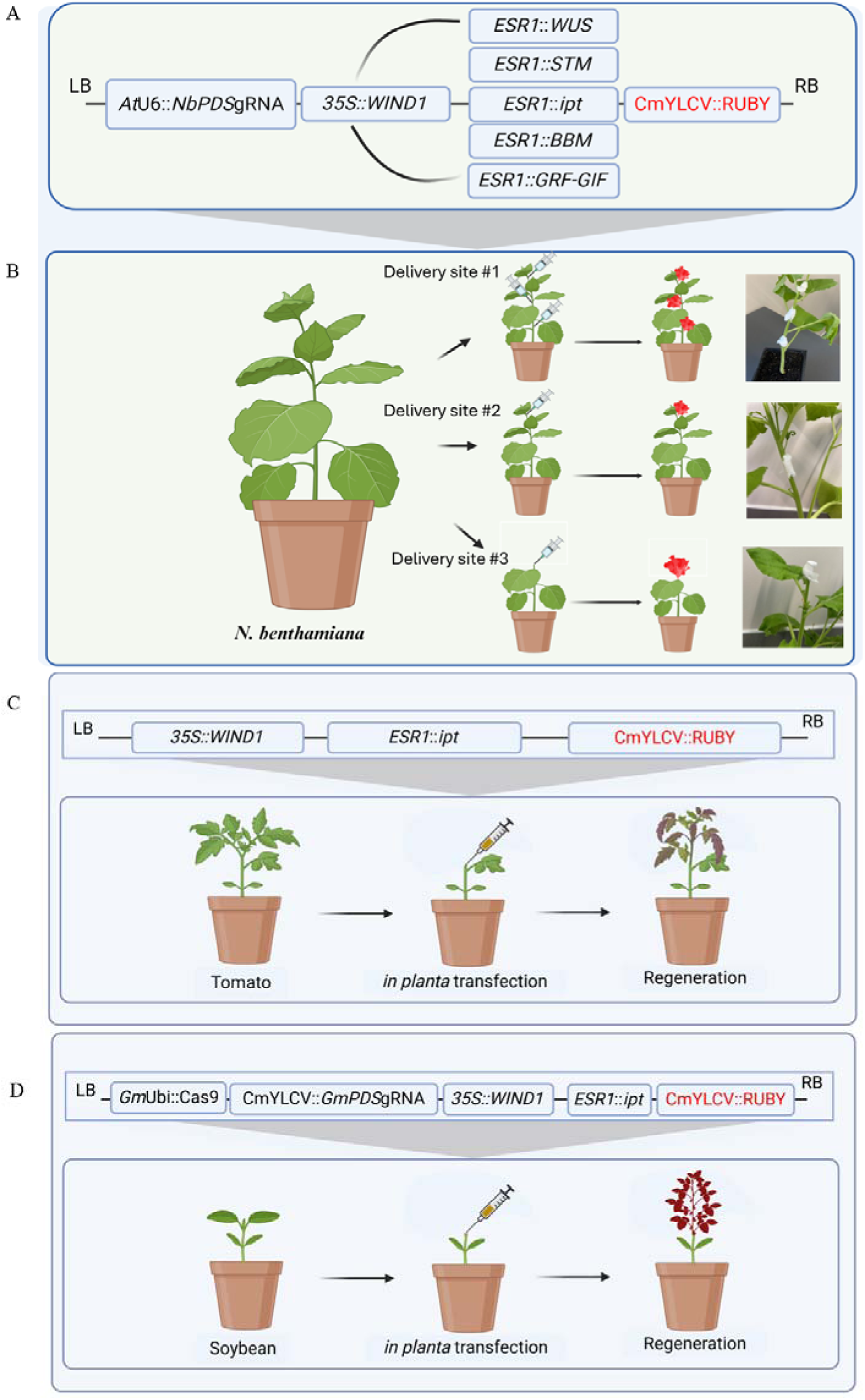
Schematic diagram showing the constructs and *in planta* transfection strategy. (**A**) Construct showing a guide RNA targeting the tobacco *PDS* gene driven by Arabidopsis Ubiquitin 6 (*At*U6) promoter, *AtWIND1* driven by the CaMV 35S promoter, An *AtESR1* promoter driving different developmental regulators (one DR in each construct), and a visible reporter gene, *RUBY* driven by the CmYLCV promoter within the T-DNA border. (**B**), Three different *in planta* transformation strategies in tobacco. After testing three different transfection strategies in tobacco, only third strategy led to *de novo* organogenesis. Therefore, we employed only a third strategy in tomato (**C)** and soybean **(D).** (**C**), Construct for *in planta* transfection and regeneration strategy in tomato. (**D**), Construct for *in planta* transfection and regeneration strategy in soybean.

**Figure 3:**
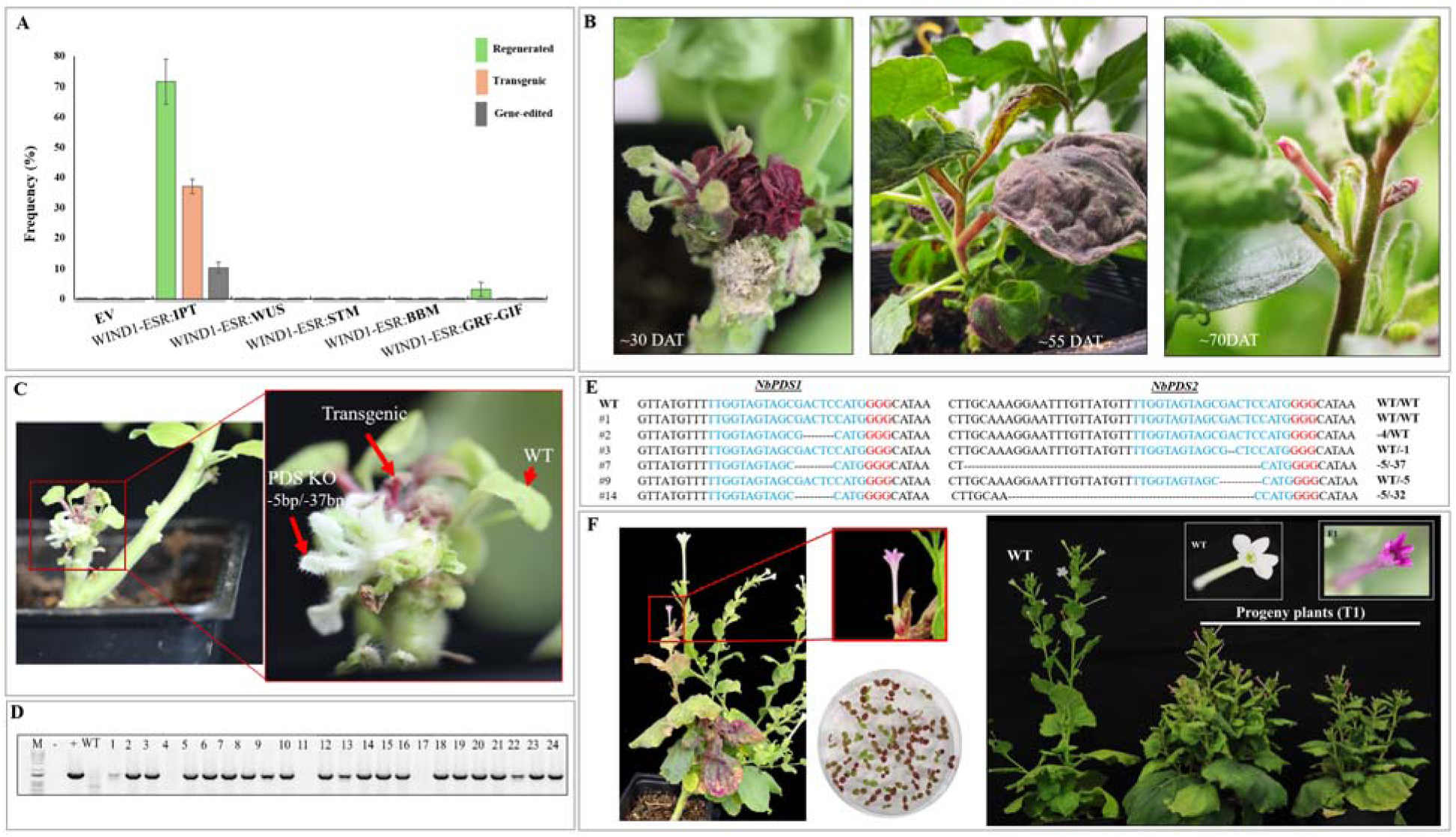
**(A)** Efficiencies of various constructs for de novo shoot induction, regeneration, transgenesis, and gene editing. **(B)** Stages of de novo shoot regeneration: Left, initiation of transgenic shoots expressing RUBY; Center, transgenic branches; Right, transgenic flowers expressing the RUBY gene. **(C)** Regeneration outcomes showing transgenic shoots (red), biallelic mutations in the PDS gene (white), and non-transgenic shoots (green). **(D)** Gel image showing PCR analysis and confirmation of the presence of the RUBY gene in progeny plants. **(E)** DNA sequence analysis for a subset of edited NbPDS alleles at the target sites. The single guide RNA (sgRNA) target region (in blue text), the PAM site (red text), and deleted nucleotide nucleotides (dashes) are indicated. **(F)** Transmission of transgenes to the next generation: Left, a representative plant showing RUBY expression in red flowers. Center (top) the zoomed in transgenic flower in T0 generation, center (bottom), segregation of transgenic (red seedlings) and non-transgenic (green seedlings) progeny germinated on ½ MS media. Right, transgenic and wild-type (WT) plants grown to maturity in pots show fading RUBY expression in late developmental stages, although flowers of transgenic plants retained a red phenotype compared to the white flowers in WT plants.

It has been well-studied that ectopic expression of *ipt* gene from a tumor-inducing (Ti) plasmid of *Agrobacterium tumefaciens* is sufficient to induce shoot primordia in dicots in *in vitro* conditions [11, 15, 36]. Notably, we observed that expression of DRs (especially *ipt*) regulated by *WIND1-ESR1* showed formation of callus followed by *de novo* shoot formations via indirect organogenesis, whereas previous studies [6, 13] showed that expression of DRs under constitutive promoters resulted in direct organogenesis with pleiotropic effects. Since the indirect organogenesis pathway involves formation of calli, the rate of *de novo* shoot induction in the present study was more uniform due to a rapid but short callus phase, resulting in ∼71% shoots regeneration between 12-18 days **(Fig. 3A**). Hence, the *AtWIND-ESR1-ipt* (*WEipt*) combination was used for downstream investigations and testing for *in planta* transformation in tomato (a close relative of tobacco) and soybean (distantly related species to tobacco).

We noted that despite stable integration of the *RUBY* gene (confirmed by PCR), the betalain accumulation (red pigmentation) was not always apparent in developing shoots, and in most cases, the red color was faint in transgenic green shoots (**Fig. 3B** and **Supplementary Fig. 3** and 4). For instance, five days after transformation, most injection sites displayed red foci until shoot emergence. However, this red coloration faded during shoot elongation, and it was variegated in mature leaves, but flowers retained bright red color (**Fig. 3B and F**). Similarly, in the subsequent generations, young tobacco seedlings germinated on petri dishes remained red (**Fig. 3F**) but faded over time (**Supplementary Fig. 4**). We also noticed that intense red pigmentation was occasionally linked to necrosis, impacting plant survival or the development of fruits (in the case of tomato; discussed in sections below). [37] have shown that despite stable integration of the *RUBY* gene in maize, some transgenic seeds did not accumulate the apparent phenotype. Similarly, [38] showed that betalain accumulation patterns were highly variable across different tissues and developmental stages of maize, but heavily pigmented plants showed stunted growth. Nevertheless, we conducted genotyping on all regenerated shoots to validate transgene insertion. Two weeks after shoot initiation, irrespective of *RUBY* expression, tissue samples were collected from each shoot to confirm transgenesis and assess gene editing outcomes (**Supplementary Fig. 5**). Genotyping (PCR analysis) revealed that 35% of the total regenerated shoots were transgenic, a notable improvement over previously reported transgenesis rates of 5-10% using single or combined DRs, including *ipt* [6, 13].

In parallel, the efficacy of the wound responsive synthetic cascade on gene-editing was evaluated. About 10% of *de novo* shoots exhibited mutations in either or both *PDS* homologs, producing mono and biallelic mutations often accompanied by semi-albino and complete albino phenotypes (**Fig. 3C and 3E, Supplementary Fig. 6**). Some transgenic (*RUBY* red) shoots showed mutations in the *PDS* gene. However, the albino phenotype was not visible due to the masking effect of red coloration from *RUBY* expression. Notably, we identified a subset of albino shoots in the T0 generation, indicating biallelic mutations (-5/-37) in both *PDS* homologs in tobacco (**Noted as plant #7 in Fig. 3C and plant #14 in Supplementary Fig. 7**D). This observation suggests that the co-expression of *WEipt1* and gene-editing reagents was effective enough to knock out *PDS* genes and induce meristematic cells simultaneously. Based on the fragment analysis and deep amplicon sequencing a range of 1 – 37 bp deletions were observed in *PDS* genes (**Fig. 3E**).

### Assessment of pleotropic effects

In this study, we also hypothesized that overexpression of *AtWIND1* in tobacco would lead to pleiotropic effects on regenerated plants similar to previous studies which used DR genes for regeneration. As expected, we observed a few plants with developmental defects, including short stature and profuse branching, and bushy leaves (**Supplementary Fig. 7**). However, compared to previous studies [6, 14, 39], frequency of pleiotropic effects in our method was less severe and only about 4% of the plants in the T0 generations showed unusual phenotypes. Furthermore, tobacco plants in the T1 generation remained relatively normal except for short stature but exhibited higher shoot biomass and flower settings than those of the wild type (**Supplementary Fig. 8A)**. Notably, the seed yield per plant in the T1 generation was more than 54% in the plants expressing *WEipt1* constructs. However, in subsequent generations, normal phenotypes were restored (**Supplementary Fig. 8B**). It has been shown that *AtWIND1* overexpression in Arabidopsis has relatively milder effects on the plant morphology, and transcriptome profiling indicated an upregulation of auxin and cytokinin biosynthesis genes and ectopic formation of tracheary elements [40]. This suggests that overexpression of *AtWIND1* in tobacco (this study) has milder or possibly positive effect compared to Arabidopsis. However, further validation experiments are required to understand the activity and regulation of *WIND1* at the molecular, physiological and morphological levels. Likewise, it has been shown that overexpression of cytokinin biosynthesis genes (tRNA-*IPT*, ADP-*IPT* or ATP-*IPT*s) significantly alters plant morphology, enhances yield and reduces the senescence [41–43]. In the present study, the pleiotropic effects of expressing *ipt* under the control of *AtWIND/AtESR*1 (*WEipt1*) cascade in tobacco were lower than those observed when expressing under constitutive or strong promoters [6, 13, 14]. In summary, controlled expression of DRs using *AtWIND1* resulted in fewer abnormal phenotypes in tobacco, tomato and soybean (as detailed below), demonstrating a promising strategy for enhancing regeneration efficiency while reduced adverse effects.

### *WEipt* mediated *in planta* transformation in tomato and soybean

The transformation of *WEipt1* with shoot at site method proved to be an efficient and seamless approach, resulting in higher regeneration, transformation, and gene-editing efficiencies in tobacco. To assess the broader applicability of this strategy, these reagents were systematically tested in crop species, including tomato and soybean. In general, conventional tissue culture-based transformation in tomato and soybean requires 4-5 and 7-9 months, respectively [44, 45]. To streamline this process, we adopted a strategy similar to the tobacco experiments (**Fig. 2**).

Tomato plants were pruned after three weeks and inoculated with *Agrobacterium* strain GV3101, carrying constructs with or without *WEipt*. As expected, injection with constructs lacking *WEipt* did not result in callus or *de novo* meristem formation. In contrast, rapid callus formation and shoot emergence (red and green shoots) were observed at injection sites treated with *WEipt* constructs (**Fig. 4A, and Supplementary Fig. 9**). For tomato, gene-editing was not evaluated since no tomato gene specific gRNA and Cas9 constructs were incorporated in *WEipt* transformation vector. Following transformation, within 3 weeks, about 53% of injection sites developed *de novo* shoots. Among these regenerated shoots, 39% were confirmed to be transgene-positive through PCR analysis resulting in overall 21% transformation efficiency (**Fig. 4B and C**). Based on findings from the tobacco experiments, we hypothesize that the remaining 62% of non-transgenic shoots might result from the transient activity of *WEipt*, the strong influence of apical dominance at the injection site, or the absence of selection pressure during these experiments. Lian et al., (2022) demonstrated the development of transgenic shoots in 13% of tomato explants injected with the *AtPLT5* gene alone. However, the constitutive overexpression of only *AtWIND1* failed to develop *de novo* shoots. This finding suggests that while *AtWIND1* activity promotes callus formation, the subsequent activity of a potent DR gene, such as *ipt*, is essential to induce de-differentiation and drive the *de novo* shoot formation in plants **(Fig. 6**). This evidence underscores the synergistic role of DRs in orchestrating differentiation and de-differentiation processes, ultimately promoting effective shoot regeneration. It also highlights the importance of combining *AtWIND1/ESR1* with complementary DRs to achieve robust and reproducible outcomes in plant transformation and regeneration. Similar to tobacco, few (∼1.5%) regenerated tomato plants showed pleotropic effects (mainly a bushy phenotype) in the T0 generation (**Supplementary Fig. 10**).

**Figure 4:**
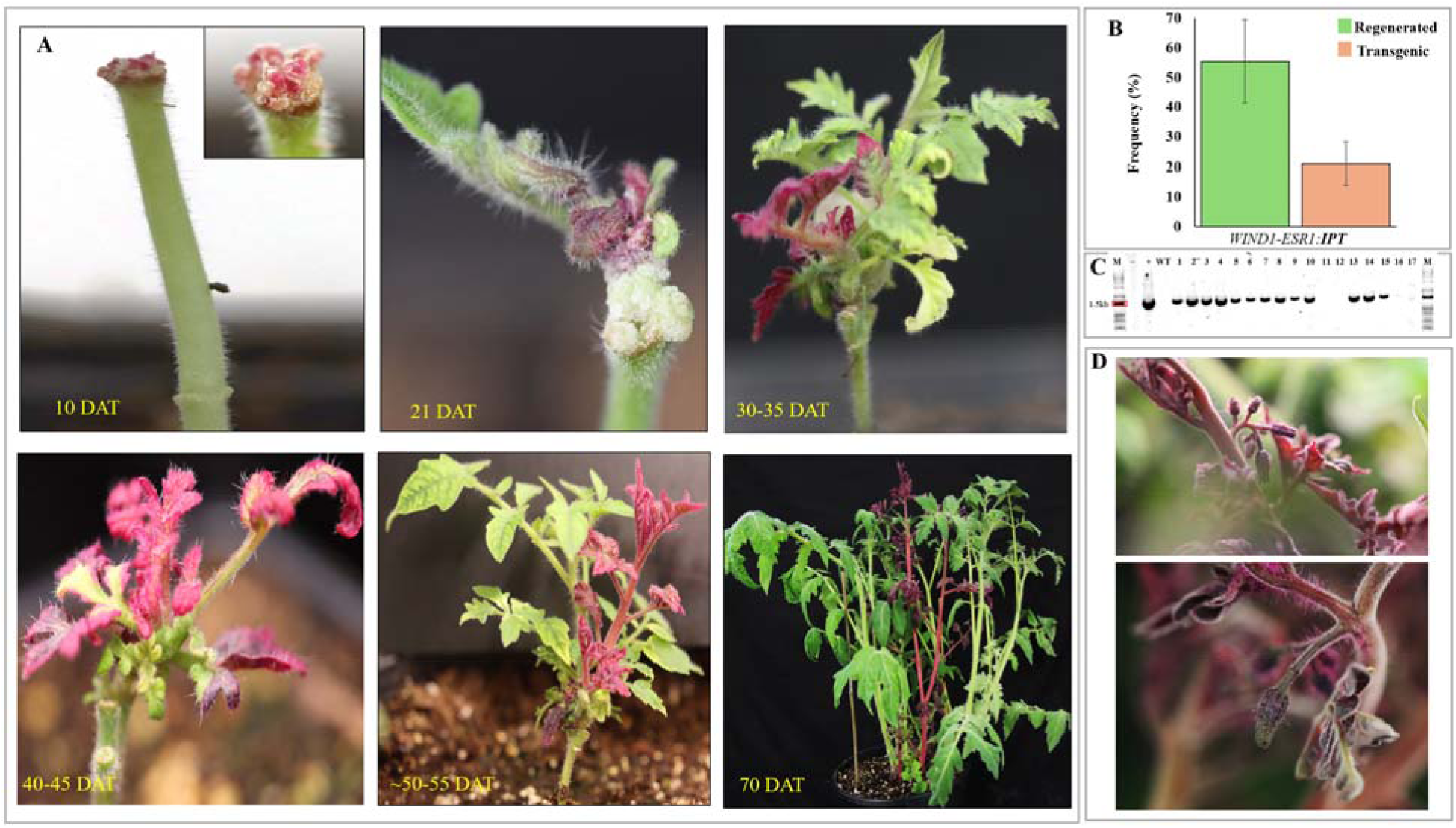
*In planta* transformation and *de novo* shoot regeneration in tomato. **(A)** Stages of *de novo* shoot regeneration in tomato, showing representative transgenic shoots (red tissues) emerging from wound sites transformed with the WEipt construct. **(B)** Quantification of transgenic frequencies in *de novo* regenerated shoots, highlighting transformation efficiency. **(C)** Gel image showing PCR analysis and confirmation of the transgene presence in representative T0 plants. **(D)** Flowers from transgenic red shoots with high betalain accumulation exhibiting impaired fruit development.

While this method successfully regenerated *de novo* shoots that produced otherwise morphologically normal tomato flowers expressing betalain, they failed to develop into fruits (**Fig. 4D**) and therefore, transmission of transgenesis to the next generation was not successful. A similar observation was reported by Lee et al., (2023), where they noted that excessive accumulation of betalain adversely impacts plant growth and development and proposed using weaker constitutive promoters as a strategy to mitigate growth retardation [38]. In the present study, over-expression of *RUBY* gene might have negatively affected fertilization or related processes in tomato flowers, however a mechanistic understanding of this observation requires additional experiments.

To further demonstrate the utility and versatility of wounding pathway-induced *de novo* shoot formation, we next targeted soybean, which is relatively more recalcitrant for genetic transformation and regeneration. In this experiment 5d old soybean seedlings were germinated in soil and subsequently used for *in planta* transformation. Similar to tobacco and tomato, soybean shoots were pruned and used for injecting *WEipt2* (that included CRISPR reagents targeting the soybean *PDS* gene) and EV construct. The injection sites were covered with cotton plugs as was done in the tobacco and tomato experiments. As expected, we observed accelerated callus formation in 97% of shoots within 5-10 days after transformation with the *WEipt2* construct. The injection sites also showed the accumulation of red foci in developing calli (**Supplementary Fig. 11**). However, despite several attempts, these plants with proliferated calli failed to develop *de novo* shoots and the callus remained compact and hard, suggesting more recalcitrant injection sites in soybean compared to tomato and tobacco. Previous soybean transformation studies have relied on laborious plant tissue culture methods and primarily using half seed [44], hypocotyl [43], and embryogenic axis [46] explants for transformation. We tested *WEipt2* using embryogenic axis explants due to the ease of explant preparation and the potential to develop a semi-tissue-culture-free transformation method in soybean (**Fig. 5**). This approach involved removing the apical meristem of the embryogenic axis to prevent the growth of untransformed primary shoots or leaf primordia (**Fig. 5A**). These explants were subsequently subjected to co-cultivation. Approximately five days after co-cultivation, 100% of the embryogenic axes displayed red pigment accumulation, while those transformed with *WEipt2* exhibited rapid callus initiation without growth hormones (**Fig. 5A, SEM image**). In 28-30 days after transformation, when explants were maintained on Gamborg B5 basal media without selection agents and hormones, initiation of green and red *de novo* shoots (81%) from the injection sites were observed **(Fig. 5B**). Since the root primordia of the explants remained intact, explants with well-established non-transgenic roots and transgenic shoots were easily transferred to soil between 36–45 days after transformation and grown to maturity. Within approximately 120 days, transgenic plants matured, producing flowers and seeds in some transgenic lines **(Fig. 5C** and **5D**). Although the regeneration frequency exceeded 80%, approximately 28% of the regenerated shoots were confirmed as transgenic via PCR, with most of these showing betalain accumulation. Similar to observations in tobacco and tomato, soybean flowers with high betalain accumulation **(Fig. 5C**) either failed to develop pods or experienced significant delays in pod formation.

**Figure 5:**
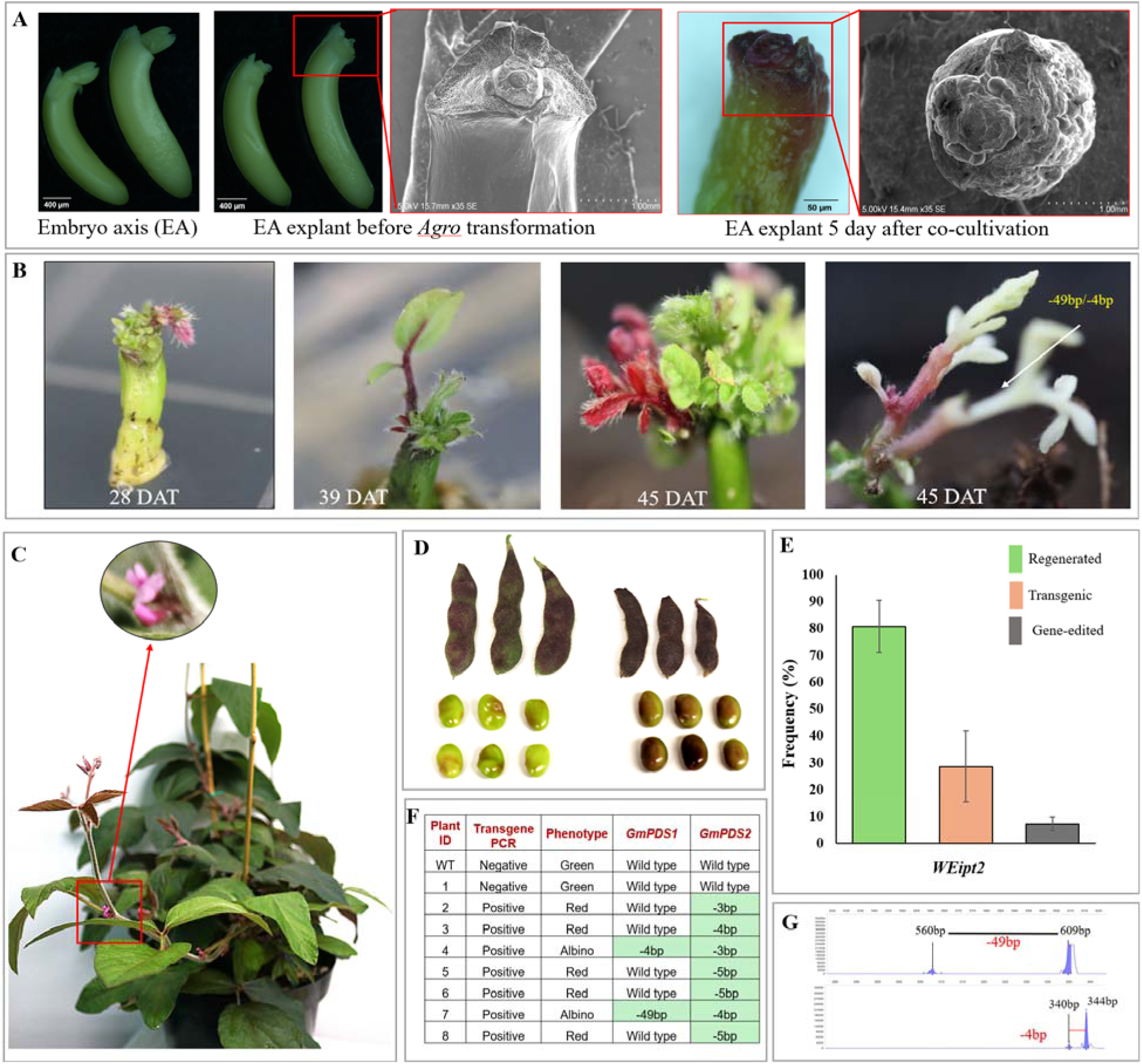
Semi-tissue culture free transformation in soybean. (**A)** Images showing embryo axis (EA) explants used for soybean transformation. Scanning electron microscopy images show the formation of callus after 3 days of transformation. **(B)** Various stages of *de novo* shoot formation, expression of the *RUBY* gene, and gene-edited shoots (albino shoots) in EA explants transformed with the *WEipt2* construct. **(C)** 40-45 days after transformation, transgenic shoots were moved to soil and continued to grow until maturity. Note the visible *RUBY* expression in young leaves and zoomed red flower above. **(D)** Pods and seeds of transgenic plants expressing *RUBY* in the early stage of pod development, whereas *RUBY* expression faded in mature pods and seeds. **(E)** Transgenesis and gene-editing frequencies in *de novo* regenerated shoots. **(F)** The table shows the T0 plant phenotypes and the corresponding editing of (*GmPDS*) genes, detected using capillary gel-based electrophoresis. **(G)** Fragment analysis showing target-specific mutation in representative plant #7, with 49bp and 4bp deletions in *GmPDS1* and *GmPDS2,* respectively.

Nevertheless, two transgenic lines successfully set pods, with pods exhibiting completed or partial red pigmentation during early development that later turned green before final maturation **(Fig. 5 C** and **5D).** The gene-editing efficiency achieved in soybean was 7.3%, wherein two plants (#4 and #7) exhibited a biallelic mutation in the *PDS* gene in T0 regenerating shoots without *RUBY* expression (**Fig. 5B, 5F** and **5G**). The remaining edited shoots displayed monoallelic mutations of *PDS* (**Fig. 5F**). This method demonstrates a promising approach for achieving transgenesis and gene editing in soybean using *WEipt2* with minimal reliance on traditional tissue culture processes.

## DISCUSSION

Regeneration is a broad term commonly used in biology that refers to a highly complex ‘reparative and restorative’ process. Reparative signifies the repair of local wounding, while restorative indicates the reconstruction of a tissue, organ, or whole individual [47]. In simple terms, regeneration provides the ability to heal from injury, survive, and initiate new life from a previous state and is the most crucial process of multicellular organisms to overcome harsh environments, diseases, and physical injury [40]. The regeneration process generally hinges on responses to growth hormones, tissue repair after wounding, or environmental cues. But importantly, these triggers collaborate with DR genes to initiate distinct responses that facilitate differentiation, dedifferentiation, and regeneration [5, 48] following the identification of pathway and molecular components downstream of different plant hormones [5, 49], researchers significantly improved methods to accelerate plant regeneration, both through traditional tissue culture [14, 18] and emerging *in planta* techniques [6, 16, 50]. While tissue culture remains the dominant approach for regenerating transgenic or gene-edited plants, it is time-intensive and laborious. In contrast, *in-planta* transformation provides a rapid alternative, bypassing tissue culture and accelerating *de novo* regeneration for transgenic and gene-edited plant production. In this study, taking advantage of *WIND1-ESR1* genes involved in the wounding pathway, we designed a synthetic cascade that enhances tissue differentiation via the wounding pathway and promotes *de novo* transgenic and gene-edited shoots by activating specific DR genes (*ipt*) without significant pleiotropic effects.

*WIND1* acts as a central regulator in promoting wound-induced cellular reprogramming in plants via systemin-independent local wound signal from *REGENERATION FACTOR1 (REF1)* and *PERP1/2 ORTHOLOG RECEPTOR-KINASE1 (PORK1)*, that regulates defense and regeneration in response [21, 27, 51]. Following the activation, *WIND1* initiate a complex cascade of molecular events, regulating downstream genes, such as *ESR1,* guiding somatic cells toward a dedifferentiated state {**Fig. 1C**; **Supplementary Fig. 1** [27, 32, 40]}. By taking advantage of *WIND1* in cellular differentiation and activating *ESR1* (**Fig. 1**), we tested the suite of DR genes that are known to play a role in organogenesis and different developmental processes in plants.

For example, *WUS* gene plays a central role in the maintenance of stem cell populations [52]; *STM* is required for maintaining undifferentiated cells [53]; *Agrobacterium ipt* produces isopentenyl AMP, a precursor of cytokinin production [11]; *BBM* promotes cell proliferation and embryo development [39]; and *GRF-GIF* complex promotes organ development [54–56]. These various combinations were delivered to three different sites in tobacco plants (**Fig. 2**), wherein delivery at pruned site resulted in successful development of *de novo* shoots. Researchers have explored diverse strategies for delivering DR genes using *in planta* transformation, each exhibiting varying efficiencies in regeneration and transgenesis [6, 13, 33]. Maher et al., (2020) implemented a technique involving the introduction of DR genes into 63–66-day-old mature tobacco plants by selectively removing all visible shoot meristems, preserving only 2-3 nodes and supporting leaves. This approach facilitated the development of shoot-like outgrowths 38-48 days post-*Agrobacterium* transfection. Similarly, Lian et al., 2022, utilized snapdragons, cultivating them for 70 days and removing primary and axillary stems at flower bud initiation to prepare for *Agrobacterium* transfection and transgenic shoots regeneration. Whereas Liu et al. (2024) developed a viral delivery system that introduces DR genes into latent axillary meristematic cells and leaves, enabling heritable gene editing in tomato and tobacco plants by inducing *de novo* shoot formation and increasing the susceptibility of meristematic cells to viral vector infection. While the above-mentioned methodologies induce *de novo* shoots bypassing tissue culture, they require extended durations (mature plants), meticulous pruning to activate axillary meristems, or use viral vector delivery systems in very specific meristematic cells.

Additionally, in these methods, initial regenerating shoots were often non-transgenic and required removal during the first 20 days post-transformation [6]. Notably, delivering constructs at pruned apical shoots (shoot at site) method significantly reduced the time required for preparing plants for transformation as well as recovering transgenic/gene-edited shoots (**Fig. 2B**). While the incorporation of *RUBY* gene along with DR genes and gene-editing reagents facilitated a non-invasive and quick assessment of transgenic shoots. Inconsistent accumulation of betalain pigments was observed in transgene positive plants and in agreement with previous reports [37], we also observed that in some cases high accumulation of betalain negatively affected plant development and seed setting. (**Fig. 2D, Supplementary Fig. 7** and 10).

Nevertheless, the transformation efficiencies achieved were 35% in tobacco, 21% in tomato, and 28% in soybean, demonstrating significant improvements compared to other *in planta* transformation methods. Additionally, successful gene-editing was achieved in tobacco and soybean wherein several *de novo* shoots showed biallelic mutation in T0 shoots. Among various DRs tested, we observed synergistic and complementary effect of *WEipt* in improving *de novo* shoot regeneration in tobacco, tomato and soybean. During this process, *WIND1* acting as a cellular reprogramming factor primarily facilitated the process of differentiation whereas *ipt* played a crucial role in indirect organogenesis via cytokinin biosynthesis [11, 57]. This observation suggests that while *WIND1* can initiate differentiation, it does not promote the maintenance or growth of plant tissue in a developmental context and hence coordinated interaction of *ipt* is required for *de novo* shoot induction and regeneration (**Fig. 6**).

**Figure 6:**
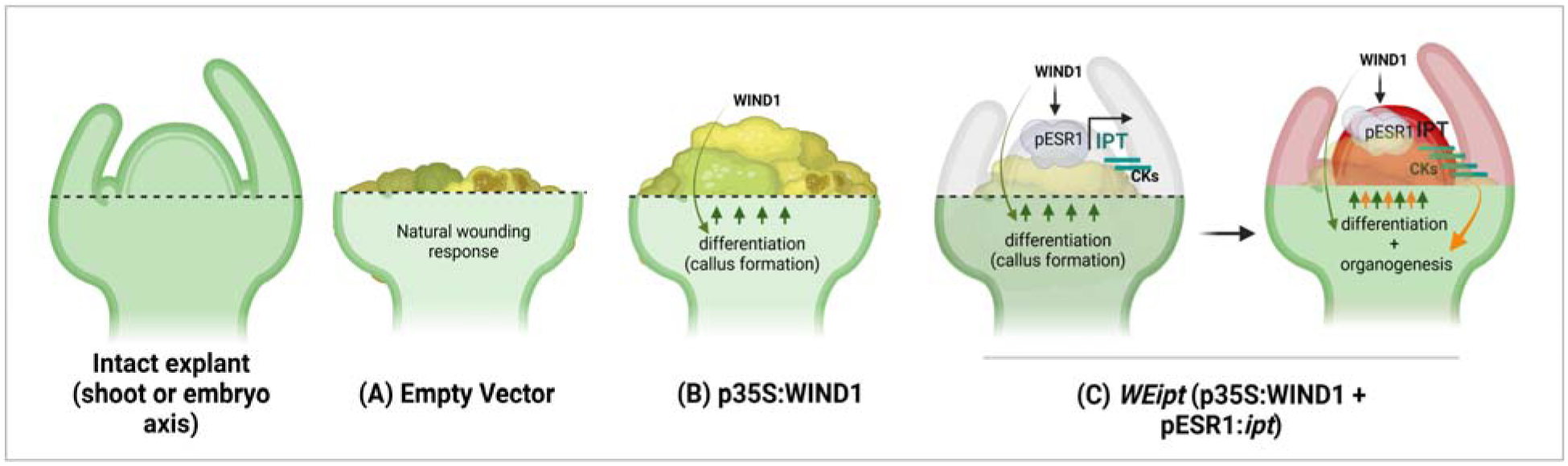
Schematic model illustrating *de novo* shoot regeneration (shoot at site). Intact shoot or embryo axis shown as an explant. The dotted line indicates the pruning of the shoot apical meristem. **(A)** Transformation with an empty vector (no DR gene) at the pruned site triggers wound healing, leading to callus formation. **(B)** Callus formation is enhanced by the activity of the *WIND1* gene, which promotes cellular reprogramming. **(C)** *De novo* shoot initiation is driven by the synergistic action of *WIND1* and *ipt* (*WEipt*). The *WIND1* reprograms somatic cells to initiate differentiation (callus formation) at the injection site, while simultaneously activating *ipt* and producing cytokinin (CKs) under the control of the ESR1 promoter. This coordinated expression accelerates differentiation through *WIND1*, while *ipt* facilitates the transition from differentiated cells to organogenesis, resulting in the formation of *de novo* shoot primordia (highlighted in red). Green arrow indicates cellular differentiation; Orange arrow indicates organogenesis.

Transcription factors and DRs genes in multicellular organisms are often expressed with precise spatial and temporal specificity, appearing in specific tissues or cell types at defined developmental stages [39]. These proteins orchestrate complex networks of gene expression, influencing multiple downstream targets that govern key processes such as growth, differentiation, and morphogenesis. While acting as primary drivers of developmental timing and spatial organization, even minor changes or deviations in their expression (compared to their normal state) can significantly impact plant morphology and physiology. For example, DR genes such as *Clavata3 (Clv3)*, *BBM*, *WUS*, and *STM* play critical roles in initiating *de novo* tissue regeneration, shifting floral meristem identity, or inducing the transition from vegetative to embryonic growth [14, 39, 58]. However, ectopic or altered expression of these genes can also lead to pronounced abnormalities in plant development, underscoring the need for precise control over their expression to avoid unintended developmental phenotypes. Lowe et al. (2016) demonstrated that ectopic expression of *BBM* and *WUS2* genes enhances regeneration in recalcitrant monocot species. However, constitutive expression of these genes also resulted in pleiotropic effects in transgenic plants, necessitating CRE-mediated excision of *BBM* and *WUS2* to achieve normal transgenic plants. Similarly, Maher et al. (2020) observed that constitutive overexpression of various DR genes resulted in a significant proportion (25-41%) of abnormal, shoot-like structures across different DR combinations. Lian et al., (2022) reported similar outcomes, with 50% abnormal morphology in regenerated transgenic plants expressing *WUS* and 22% abnormalities in those expressing *PLT5* as a developmental regulator. In another significant study, Debernardi et al. (2020) showed that although constitutive expression of the *GRF-GIF* chimera markedly improved transformation and regeneration efficiencies in wheat, it also led to 24% decrease in grain yield per spikelet and 13.7% increase in grain weight. Collectively, these findings underscore that deviations from normal DR gene expression may disrupt the plant growth and yield traits and therefore carefully controlled expression strategies are necessary to overcome the unintended phenotypic consequences. Interestingly, our study showed distinct response of *WIND1-ESR1:ipt* combination on pleiotropic effect as compared to constitutive overexpression of other DR genes in *in vitro* regeneration of maize, sorghum and sugarcane [14, 59] and *in planta* transformation of tobacco, potato [6], snapdragon, Bok choy and tomato [13]. Our approach showed minimum (4%) to no pleiotropic effect of ectopic expression of *WEipt* on plant growth and development in T0 and subsequent generation of tobacco, tomato and soybean (**Fig. 3 – 5**). Although, understanding the minor or reduced pleiotropic effect in our study necessitates further investigation, we hypothesized that the influence of *WEipt* remained localized to shoot regeneration event rather than broader growth patterns. Ectopic expression of *WIND1* at injection site reprogrammed somatic cells to initiate differentiation but simultaneously (via *ESR1*), it activated *ipt* gene allowing transition of differentiated cells to a pluripotent state, allowing for new *de novo* shoot formation (**Fig. 6**).

In conclusion, this study highlights the synergistic action and transformative potential of the *WEipt* to accelerate *de novo* shoot regeneration with minimal pleiotropic effects. By activating a wound-induced cellular reprogramming cascade, *WIND1* facilitated somatic cell differentiation, while *ipt* mediated cytokinin biosynthesis, enabling a seamless transition to pluripotent states and subsequent shoot formation (**Fig. 6**). Importantly, co-expression of gene-editing reagents with *WEipt* facilitated precise targeted mutations, establishing a scalable and efficient platform for crop improvement. The indirect organogenesis pathway improved regeneration rates and also provided a uniform and rapid timeline for shoot induction, overcoming the variability and inefficiencies observed in earlier methodologies. Compared to previous approaches relying on constitutive overexpression of DR genes, which often result in developmental abnormalities, the targeted and localized expression of *ipt* achieved consistent and efficient regeneration across tobacco, tomato, and soybean. This approach holds a significant promise for *in planta* transformation, reducing the dependency on tissue culture and accelerating the generation of transgenic or gene-edited plants.

## MATERIALS AND METHODS

### Plant materials, and growth conditions

A stable transgenic *N. benthamiana* expressing 35S::Cas9 was kindly provided by Prof. Danial Voytas Lab, University of Minnesota. Tomato seeds (*Solanum lycopersicum var* money maker) were purchased from the marketplace and used to evaluate various constructs. Both the plant species were germinated under long-day conditions at 22□, exposed to 16 hours of white light and 8 hours of darkness at 20□. Similarly, for soybean *in planta* transformation, mature and dried soybean seeds (genotype ‘Williams 82’) were sown in soil and germinated at 25□, exposed to 16 hours of white light and 8 hours of darkness.

### Vector design and construction

All DNA constructs were created in accordance with plant genome engineering toolkit [60]. As indicated in **Supplementary Table 1**, the DNA cassettes included CRISPR reagents (Cas9 and guide RNAs), DRs, and reporter genes cloned in one of the modular vectors that are finally assembled on a T-DNA destination vector by Golden Gate cloning strategy. *RUBY* [35] as a visible reporter gene under the Cestrum Yellow Leaf Curling Virus (CmYLCV) promoter [61] was cloned into module A. Guide RNAs targeting *PDS* genes in tobacco and soybean were cloned in module B under Arabidopsis Ubiquitin 6 (U6) and CmYLCV promoters respectively. The Arabidopsis *ESR1* (AT1G12980) promoter (1046 bp) was amplified from the Arabidopsis Col-0 genotype and cloned into module C’. This module was subsequently used for cloning the DR genes. Similarly, *WIND1* (AT1G78080) from Arabidopsis Col-0 was cloned in module D under the CaMV 35S promoter [62]. For soybean transformation, Cas9 protein was cloned in Module A driven by soybean ubiquitin (*Gm*Ubi) promoter and visible reporter gene *RUBY* was cloned to module E driven by CmYLCV promoter. All the modules were finally assembled into T-DNA transformation backbone using Golden Gate Assemblies [60]. These assembled constructs were transformed into the *Agrobacterium tumefaciens* strain GV3101 to transform tobacco and tomato, and strain 18r12 for soybean transformation. Before plant transformation, confirmation of successful delivery of T-DNA carrying the *RUBY* gene was visualized through agroinfiltration of tobacco leaves (**Supplementary Fig. 12**).

### Assessment of *WIND1-ESR1* cascade

To confirm the role of *AtWIND1* in tissue differentiation, three different constructs were generated (**Supplementary Table 1,** *RUBY*-Empty, *ESRruby*, and *WEruby*). The construct *RUBY*-Empty (proCmYLCV::*RUBY*) and *ESRruby* (pro*AtESR1*::*RUBY*) were used as negative controls for callus formation and induction of *ESR1* activity, respectively. To verify if *AtWIND1* activates the *AtESR1* promoter and induces callus formation, construct WE*ruby* comprising pro35S::*WIND1* and pro*AtESR1*::*RUBY* were developed. These constructs were transformed into *Agrobacterium* strain GV3101 and tobacco explants (petioles and internodes) were used for transformation. The transformed explants were cultured in ½ strength hormone-free MS media for 4 weeks. Areas of callus induction were measured to assess the *AtWIND1* activity in callus formation.

### *In planta* transformation in tobacco, tomato, and soybean

The constructions detailed in **Supplementary Table 1** were utilized to assess *in planta* shoot formation, transgenesis and gene editing. For *in planta* transformation of tobacco, Cas9 positive transgenic seeds were germinated and grown for approximately 4 weeks and then injected with the *Agrobacterium* carrying the construct with one of the DRs at various sites that do not have meristematic activity. A single colony of *Agrobacterium* harboring synthetic cascade (*WEipt* and CRISPR reagents) was inoculated in 30mL yeast extract peptone (YEP) broth with antibiotics (Kanamycin, 50mg/L, Rifampicin, 10mg/L, and Gentamicin, 30mg/L for GV3101) and grown (shaking at 160rpm/12 hours) at 28°C. After 12 hours, the culture was pelleted by centrifuging at 4500 rpm for 10 minutes. The pellet was resuspended with the transfection buffer (10mM MES, 150µM acetosyringone, and 10mM MgCl_2_) until the OD_600_ was between 0.2-0.8 [6]. Three different wounding strategies or locations were used for *in planta* transformation. First, multiple wounding sites in the internode of the plant; second, a single wounding site at the internode; and third, pruning the plant at the internode. These sites were used to inoculate the *Agrobacterium* with respective vectors carrying DR cascades. After inoculation, the wounding area was covered with sterile cotton plug soaked in the same *Agrobacterium* solution, creating a localized microenvironment conducive to co-cultivation. Next, plants were placed in the dark in a high humidity (80%) growth chamber at 22°C for 3 days. After 3 days, the cotton plug was removed, and plants were grown in a normal condition at 16/8-hour day/night cycle at 22/20°C. Plants were routinely observed for shoot-like growth at the site of injection. For tomato, only one of the three strategies was adapted, based on the successful *N. benthamiana* experiments. The tomato plants between 3 and 4 weeks old were pruned and inoculated with *Agrobacterium* carrying *WEipt* at the wound site. The gene-editing experiment was not performed in tomato and therefore the *WEipt* construct did not harbor gene-editing reagents. The rest of the protocol followed the same as the tobacco experiment described above.

For soybean *in planta* transformation, healthy soybean seedlings were excised above the cotyledon before they formed the first true leaf. The *WEipt2* construct harboring gene-editing reagents targeting soybean *PDS* gene and *ipt* was injected following wounding, and the wound sites were plugged in with cotton plugs soaked with *Agrobacterium* solution. Plants were incubated in the dark for 3 days in high humidity (80%). After 3 days, cotton plugs were removed, and the plants were placed in a growth chamber with similar growth conditions as they were germinated. The infusion sites were observed for callus (transgenic; red or normal) formation and regeneration.

For soybean embryogenic axis transformation, mature and healthy seeds of soybean genotype Williams 82 were surface sterilized as detailed in [44]. Following imbibition, seeds were rinsed with sterile water twice and prepared for isolation of the embryogenic axis. The shoot apical meristem of the embryogenic axis was carefully removed and used for co-cultivation. The embryogenic axes were transformed by sonicating them with *Agrobacterium* (strain 18r12) solution for 30 seconds and shaking them in a rotary shaker at 75RPM for 30 minutes. The agrobacterial solution was drained by pipetting after shaking, and the explants were placed in petri dishes containing sterile filter paper and presoaked in ½ strength MS media. The plates were sealed with parafilm and placed in a growth chamber with 23°C under 16hr light/8hr dark photoperiod for 5 days. After 5 days of co-cultivation, the explants were transformed aseptically to ½ strength plant growth regulator (hormone) free Gamborg B5 basal media supplemented with antibiotics (Timentin, 50mg/L) to eliminate the excessive Agrobacterial growth. Regeneration of non-transgenic (green) and transgenic (red) shoots was observed, and the count of each was recorded.

### Phenotypic analysis of regeneration, transgenesis, and gene editing

Any shoot-like growth at the site of injection was considered as a *de novo* shoot induced by developmental regulators. The *de novo* meristems were visually observed for white (indicating possible loss of function of *PDS* by gene editing), red betalain pigment (indicating expression of the *RUBY* transgene), and green (functional *PDS* and no transgene expression).

### Genotypic analysis for gene editing and transgenesis

DNA was extracted from the leaf tissues collected from white, red, and green meristems using CTAB (cetyl trimethyl ammonium bromide) [63]. A fragment of *RUBY* gene was amplified for transgene PCR analysis. For analysis of gene-editing events in tobacco, both the copies of the *PDS* gene were amplified using forward primer ∼212bp upstream and reverse primer ∼232bp downstream of the sgRNA target site, with a total amplicon size of ∼464bp. Primary amplicons were re-amplified using a 19bp M13 tail flanked forward and a gene specific reverse primer. In a third step, final amplicons were generated by using a FAM dye-labeled M13 primer and the same gene-specific reverse primer. PCR products were diluted 1:25 fold, and 1µL mixed with 9µL HiDi formamide and 0.5µL LIZ600 size standard (Thermo Fisher Scientific, Waltham, MA, USA). The mixture was denatured at 95 for 5 minutes and immediately kept in ice. Capillary gel-based electrophoresis was performed using seqstudio genetic analyzer (Thermo Fisher Scientific, Waltham, MA, USA). Peak intensity and size were detected by microsatellite analysis (MSA) software (Thermo Fisher Scientific, Waltham, MA, USA). For the confirmation of transgenesis, a fragment of *RUBY* was amplified using primers listed in **Supplementary Table 2.** Deep amplicon sequencing and analysis was performed as detailed in [49].

## Supplementary Figure Legends

**Supplementary Figure 1:** Transcriptional regulation after wounding in plants. Wounding leads to the activation of endogenous hormone signaling pathways and wound-induced stress signal/s. WOUND-induced signals such as *REF1-POKR1* and *WIND1* are activated at the wound site and promote callus formation. *WIND1* binds to the *ESR1* promoter and transcriptionally activates the expression of *ESR1*, inducing direct shoot regeneration by regulating genes known as DRs without the supplementation of exogenous phytohormones. Wounding induces the expression of *PLT3*, *PLT5*, and *PLT7* which are required for callus formation. In auxin rich media, wounding induces callus formation through ARFs (*ARF7*, *ARF19*)-mediated activation of LBDs (*LBD16*, *LBD18*, and *LBD29*). These LBDs, in turn promote a battery of genes responsible for callus formation and proliferation. Wounding triggers the expression of LOGs (*LOG1*, *LOG4*, and *LOG5*), and *IPT3*, thereby activating the cytokinin biosynthetic pathway and inducing the expression of ARRs, which in turn induces the expression of *CYCD3* and callus formation. The fate of a callus depends on its transfer to SIM or RIM. In SIM media, the expression of CUCs leads to shoot progenitor formation, and *WUS* leads to shoot regeneration. Similarly, in RIM, LBDs and *WOX* lead to root regeneration. **Abbreviations:** *REF1*: Regeneration Factor 1, *PORK1*: *PEPR1/2* Ortholog Receptor-like Kinase1, *ESR1*: Enhancer of Shoot Regeneration 1, *PLT: PLEOTHERA*, *ARF*: Auxin Response Factor, *LBD*s: Lateral Boundary Domains, *LOG*s: Lonely Guys, *ipt*: Isopentenyl Pyrophosphate Transferase, *CYCD3*: Cyclin D3, *CUC*s: Cup-shaped cotyledons, *WUS: WUSCHEL, WOX: WUSCHEL* -related Homeobox, DRs: Developmental Regulators, SIM: Shoot Induction Media, RIM: Root Induction Media

**Supplementary Figure 2:** *in planta* transformation strategies in *N. benthamiana*. Outcomes of different *in planta* transformation strategies in tobacco (**A-C**). Transformation at multiple sites (non-meristematic internodes) leads to the healing of the wound without any signs of callus formation or regeneration (**Panel A**). Transformation at single site (non-meristematic internode) leads to callus formation but no regeneration (**Panel B**). Pruning of the plant apical meristem led to callus formation and *de novo* shoot formation (**Panel C**).

**Supplementary Figure 3:** *in planta* transformation in *N. benthamiana* leading to regeneration of *RUBY* (betalain) expressing (red) shoots in the green mother plants. The representative image shows variable expression of *RUBY* in different T0 plants.

**Supplementary Figure 4:** Transgenic shoots transmit the transgene to the progeny. The plants in the progeny (T1) were analyzed for the transmission of transgenesis (A fragment of *RUBY* was PCR-amplified from the DNA extracted from these plants). However, the expression of RUBY was observed to vary, as shown in the figure above. Based on the pattern of *RUBY* expression in the progeny plants, they were scored as follows: panel B ‘Green’, panel C as ‘variegated’, and panel D as ‘Red’ with compared to the Wild Type, panel A. The tube show, juice samples extracted from 200mg of leaf tissue in 2mL water. The intensity of betalain is shown for comparative purposes.

**Supplementary Figure 5:** PCR genotyping of representative *de novo* meristems, showing amplification of *ipt* to verify T-DNA insertion in tobacco (amplicon size 723bp). This verified the presence of T-DNA insertions in tobacco. (M= Marker; -= negative control and += positive control)

**Supplementary Figure 6:** *In planta* transformation in tobacco with *WEipt1* construct induced regeneration of semi-albino shoots resulting from mono-allelic mutations in the (*PDS*) gene.

**Supplementary Figure 7:** Examples of *in planta* transformation in tobacco with *WEipt1* construct leading to the regeneration of *de novo* meristem with developmental defects in T0 generation. (**A**); WT. (**B-D**); abnormal stature.

**Supplementary Figure 8:** Phenotypes of the progeny plants. The phenotype of T1 the plants was relatively normal except for short stature, multiple branching, and flower setting in comparison to WT **(A).** The phenotype of the plants was restored to normal in subsequent generations **(B).**

**Supplementary Figure 9:** *In planta* transformation of tomato with *WEipt* construct induced accelerated dedifferentiation and *de novo* meristem formation at the site of wounding. Transient expression of *RUBY*, five days post transfection **(A)** and ten days post-transfection **(B).** Regenerating callus with a non-transgenic (green) shoot and a transgenic callus **(C).** Transgenic callus that never regenerated **(D).** Regeneration of transgenic and non-transgenic shoots induced by *WEipt* (**E-H)**.

**Supplementary Figure 10:** Example of pleiotropic effects of *WEipt* observed in tomato in the T0 generation.

**Supplementary Figure 11:** *in planta* transformation of soybean with *WEipt2* led to rapid callus formation at the transfusion site without shoot regeneration. (Insets show images at larger scale).

**Supplementary Figure 12:** Visual confirmation of successful delivery of T-DNA carrying *RUBY* as a visible reporter gene, achieved through tobacco leaf agroinfiltration.

## Supplementary Table Legends

**Supplementary Table 1**: List of vectors used in this study.

**Supplementary Table 2**: List of gRNA and the gRNA sequence targeting different genes in different plant species.

**Supplementary Table 3**: List of primers used in this study.

## Authors Contributions

AOK: methodology, experimental design, formal analysis, data interpretation and writing original draft; KG: genotyping and data interpretation; AA: methodology; VD, VR, RMS, LHE, FZ: experiment design, manuscript editing; GBP: Conceptualization, formal analysis, funding acquisition, supervision and writing and editing manuscript.

## Acknowledgements

G.B.P is grateful to the State of Texas’ Governor’s University Research (GURI) for the research funding. A.A., V.R., and F.Z. are supported by the National Science Foundation (IOS-2040218 and IOS-2206920) awards. G.B.P., R.M.S., and F.Z. were supported by the USDA NIFA award #2021-67013-34565. We thank Dr. Danial Voytas and Dr. Colby Starker, University of Minnesota, for sharing the modular clones and Cas9 positive transgenic *N. benthamiana* seeds.

