## Supplementary Table for "Shoot at Site: Advancing *in planta* transformation, regeneration and gene-editing through a cascade of wounding-mediated developmental regulators"

**SUPPLEMENTARY TABLES**

**Supplementary Table 1**: List of vectors used in this study

| **Plasmid id** | **Module A** | **Module B** | **Module C/C'** | **Module D/D’** | **Module E** | **Vector backbone** |
| --- | --- | --- | --- | --- | --- | --- |
| **Empty Vector (EV)** | Empty | Empty | Empty | - | - | pTRANS_231 |
| ***RUBY*-Empty** | CmYLCV::*RUBY* | Empty | Empty | - | - | pTRANS_231 |
| ***ESRruby*** | Empty | Empty | *ESR1::RUBY* | Empty | - | pTRANS_231 |
| ***WEruby*** | Empty | Empty | *ESR1::RUBY* | 35S::*WIND1* | - | pTRANS_231 |
| ***WEipt*** | CmYLCV::*RUBY* | Empty | *ESR1::ipt* | 35S::*WIND1* |  | pTRANS_231 |
| ***WEwus*** | CmYLCV::*RUBY* | *NbPDS*gRNA | *ESR1::WUS* | 35S::*WIND1* | - | pTRANS_231 |
| ***WEstm*** | CmYLCV::*RUBY* | *NbPDS*gRNA | *ESR1::STM* | 35S::*WIND1* | - | pTRANS_231 |
| ***WEbbm*** | CmYLCV::*RUBY* | *NbPDS*gRNA | *ESR1::BBM* | 35S::*WIND1* | - | pTRANS_231 |
| ***WEgrf*** | CmYLCV::*RUBY* | *NbPDS*gRNA | *ESR1::GRF_GIF* | 35S::*WIND1* | - | pTRANS_231 |
| ***WEipt1*** | CmYLCV::*RUBY* | *NbPDS*gRNA | *ESR1::ipt* | 35S::*WIND1* | - | pTRANS_231 |
| ***WEipt2*** | *Gm*Ubi::Cas9 | *GmPDS*gRNA | *ESR1::ipt* | 35S::*WIND1* | CmYLCV::*RUBY* | pTRANS_231 |

**Supplementary Table 2**: List of gRNA and the gRNA sequence targeting different genes in different plant species

| S. No. | Target plant | Target gene | gRNA1/PAM | gRNA2/PAM |
| --- | --- | --- | --- | --- |
| 1 | *N. benthamiana* | *PDS* | TTGGTAGTAGCGACTCCATG/GGG | - |
| 2 | *Glycine max* | *PDS* | GCAAAATATTTGGCTGATGC/TGG | GAGAACTTGGCATTAATGAT/CGG |

**Supplementary Table 3**: List of primers used in this study

| **S. No.** | **Forward (5’-3’)** | **Reverse (5’-3’)** | **Comments** |
| --- | --- | --- | --- |
| 1 | ATGGATCATGCGACCCTCG | AGGCCCGGGGTTCTCTTC | Genotyping primers amplifying part of RUBY |
| 2 | TTTAGGTTCACAAGTGGGACA | TGATAATAACGCCGCCTCCA | Amplifying *N. benthamiana* *PDS1* and *PDS2* |
| 3 | CGCAAACGCAGCCATTACAA | TTTTGGTTTCTAGGGTTTTGGTTTG | Amplifying Arabidopsis *ESR1* promoter |
| 4 | TCTTGCGGAGCTAGCATA | CTAAGCTAGAATCGAATCCCAAT | Amplifying Arabidopsis *WIND1* CDS |
| 5 | ATGGATCATGCGACCCTCGCCATG | TCACTGGAGGCTTGGCTCAAG | Amplifying RUBY CDS |
| 6 | CTTCGTTGAACAACGGAAACTCGA | AATCACTACTTCGACTCTAGCTGT | Primers to amplify AtU6 promoter |
| 7 | ACACTCTTTCCCTACACGACGCTCTTCCGATCTTTTAGGTTCACAAGTGGGACA | GACTGGAGTTCAGACGTGTGCTCTTCCGATGATAATAACGCCGCCTCCA | Primers for deep amplicon seq with Illumina adapters for *NbPDS1* and *NbPDS2.* |
| 8 | TGTAAAACGACGGCCAGTTTTAGGTTCACAAGTGGGACA | TGATAATAACGCCGCCTCCA | M13 flanked forward and gene-specific reverse primer for *NbPDS1* and *NbPDS2* for capillary gel-based electrophoresis |
| 9 | GGCCAGGAAGAAGAAAAGCC | CAACACATGAGCGAAACCCT | Soybean gRNA amplification prime |
| 10 | TGCTCTTGTTGTTCTGGGTG | TGCTCTTGTTGTTCTGGGTG | GmPDS1a target amplification |
| 11 | GGTTGCTGCATGGAAAGACA | CATTTAATGGGGCGGGAAGG | GmPDS1b target amplification |
| 12 | TGGGCTACGCATTAGGAATTTC | AGATGTGTAGGCCTGTCTCG | GmPDS2a target amplification |
| 13 | TGAGTTAGTCCATGCCACACT | TTCAATGGGGAGGGAAGAACT | GmPDS2b target amplification |
