## Supplementary Figures for "Shoot at Site: Advancing *in planta* transformation, regeneration and gene-editing through a cascade of wounding-mediated developmental regulators"

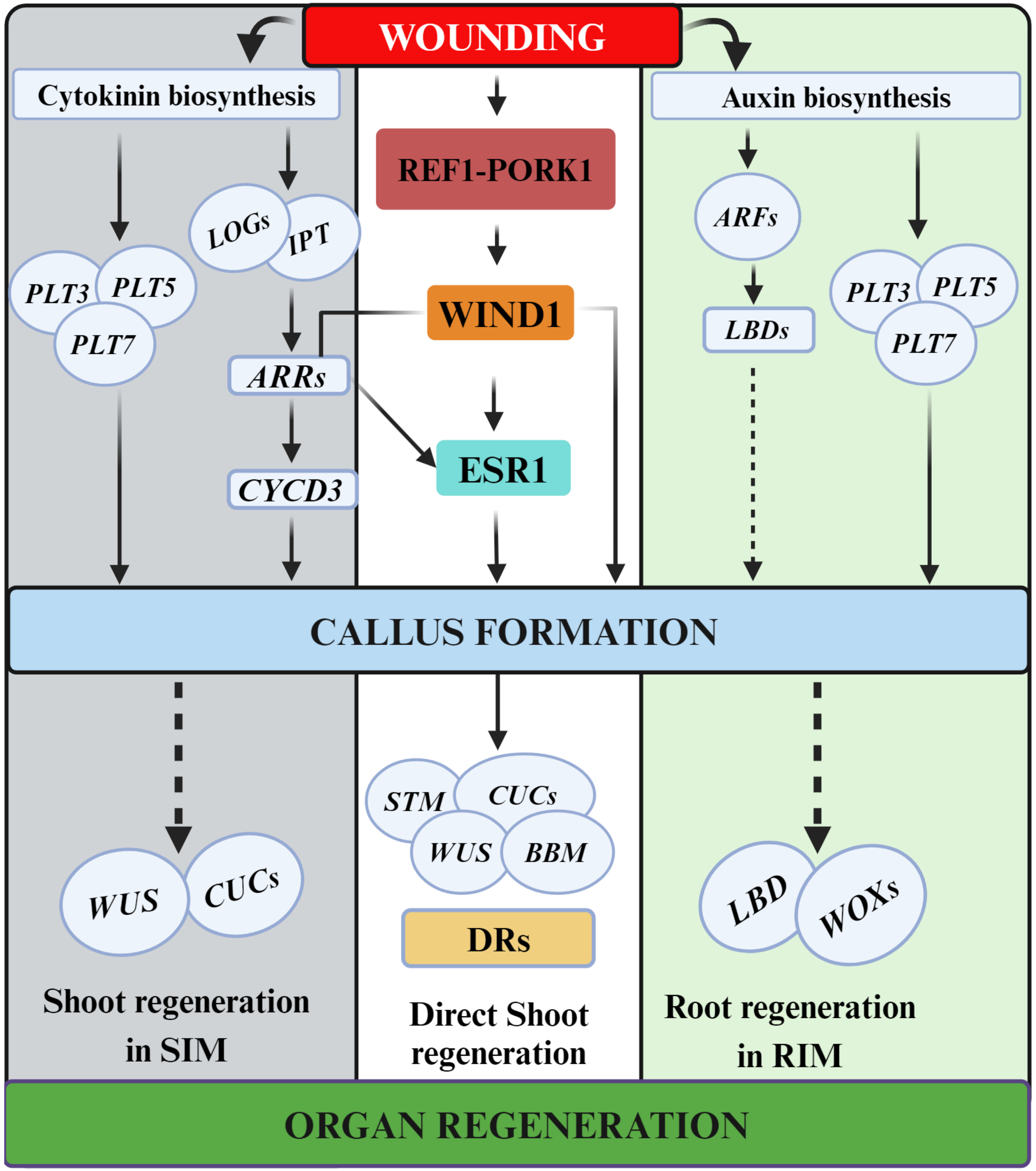


**Supplementary Fig. 1** | Transcriptional regulation after wounding in plants. Wounding leads to the activation of endogenous hormone signaling pathways and wound-induced stress signal/s. WOUND-induced signals such as *REF1-POKR1* and *WIND1* are activated at the wound site and promote callus formation. *WIND1* binds to the ESR1 promoter and transcriptionally activates the expression of *ESR1*, inducing direct shoot regeneration by regulating genes known as DRs without the supplementation of exogenous phytohormones. Wounding induces the expression of *PLT3*, *PLT5*, and *PLT7* which are required for callus formation. In auxin rich media, wounding induces callus formation through *ARF*s (*ARF7*, *ARF19*)-mediated activation of *LBD*s (*LBD16*, *LBD18*, and *LBD29*). These *LBD*s, in turn promote a battery of genes responsible for callus formation and proliferation. Wounding triggers the expression of *LOG*s *(LOG 1*, *LOG 4*, and *LOG 5*), and *IPT3*, thereby activating the cytokinin biosynthetic pathway and inducing the expression of ARRs, which in turn induces the expression of *CYCD3* and callus formation. The fate of a callus depends on its transfer to SIM or RIM. In SIM media, the expression of CUCs leads to shoot progenitor formation, and *WUS* leads to shoot regeneration. Similarly, in RIM, *LBD*s and *WOX* lead to root regeneration.

Abbreviations: REF1: Regeneration Factor 1, PORK1: PEPR1/2 Ortholog Receptor-like Kinase1, *ESR1*: *Enhancer of Shoot Regeneration 1*, *PLT: PLEOTHERA*, *ARF: Auxin Response Factor*, *LBDs: Lateral Boundary Domains*, *LOGs: Lonely Guys*, *IPT: Isopentenyl Pyrophosphate Transferase*, *CYCD3: Cyclin D3*, *CUCs: Cup-shaped cotyledons*, *WUS: WUSCHEL*, *WOX: WUSCHEL - related Homeobox*, DRs: Developmental Regulators, SIM: Shoot Induction Media, RIM: Root Induction Media

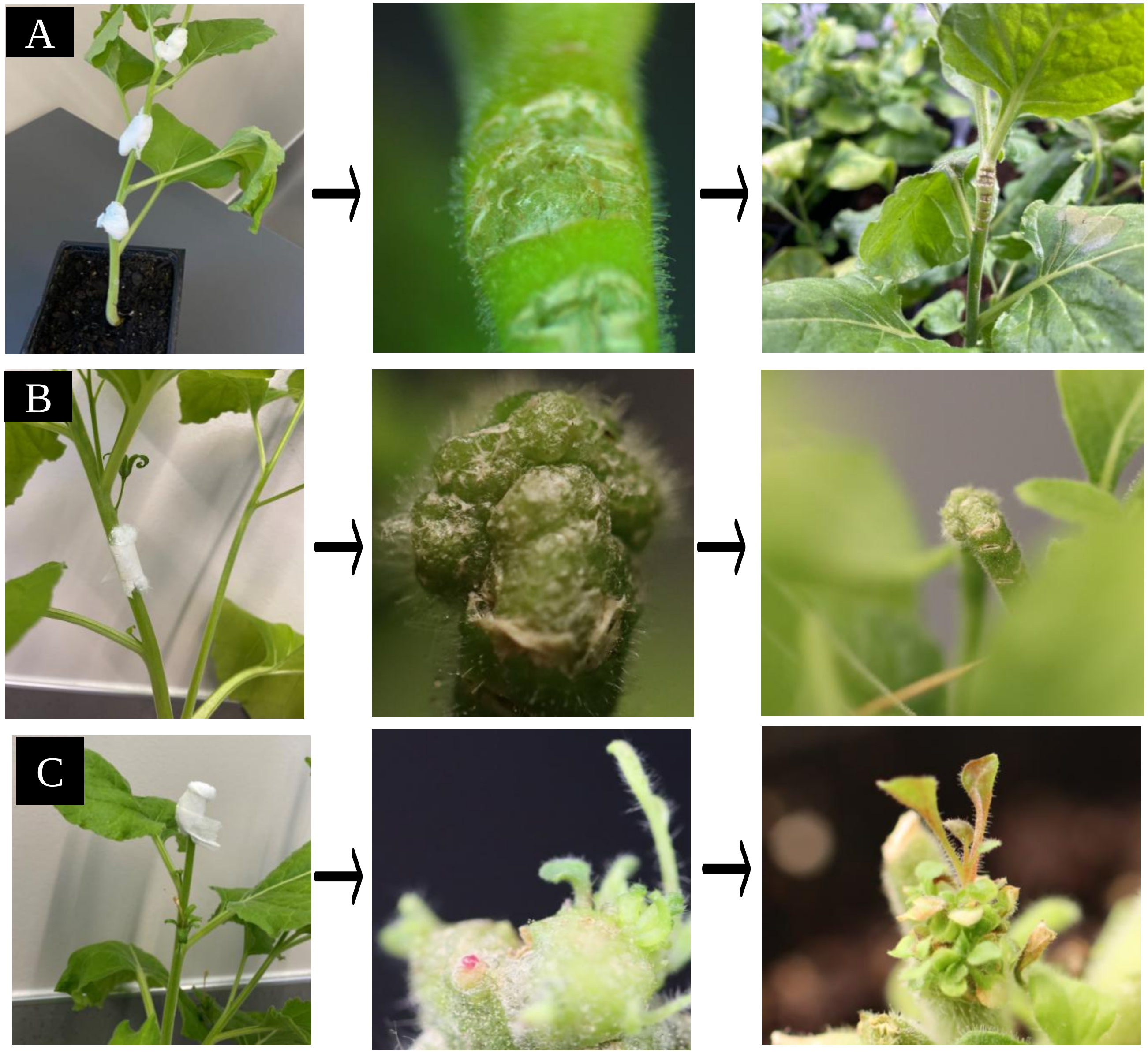


**Supplementary Fig. 2 |** *in planta* transformation strategies in *N. benthamiana*. Outcomes of different *in planta* transformation strategies in tobacco (**A-C**). Transformation at multiple sites (non-meristematic internodes) led to the healing of the wound without any signs of callus formation or regeneration (**Panel A**). Transformation at single site (non-meristematic internode) led to callus formation but no regeneration (**Panel B**). Pruning of the plant apical meristem led to callus formation and *de novo* shoot formation (**Panel C**)

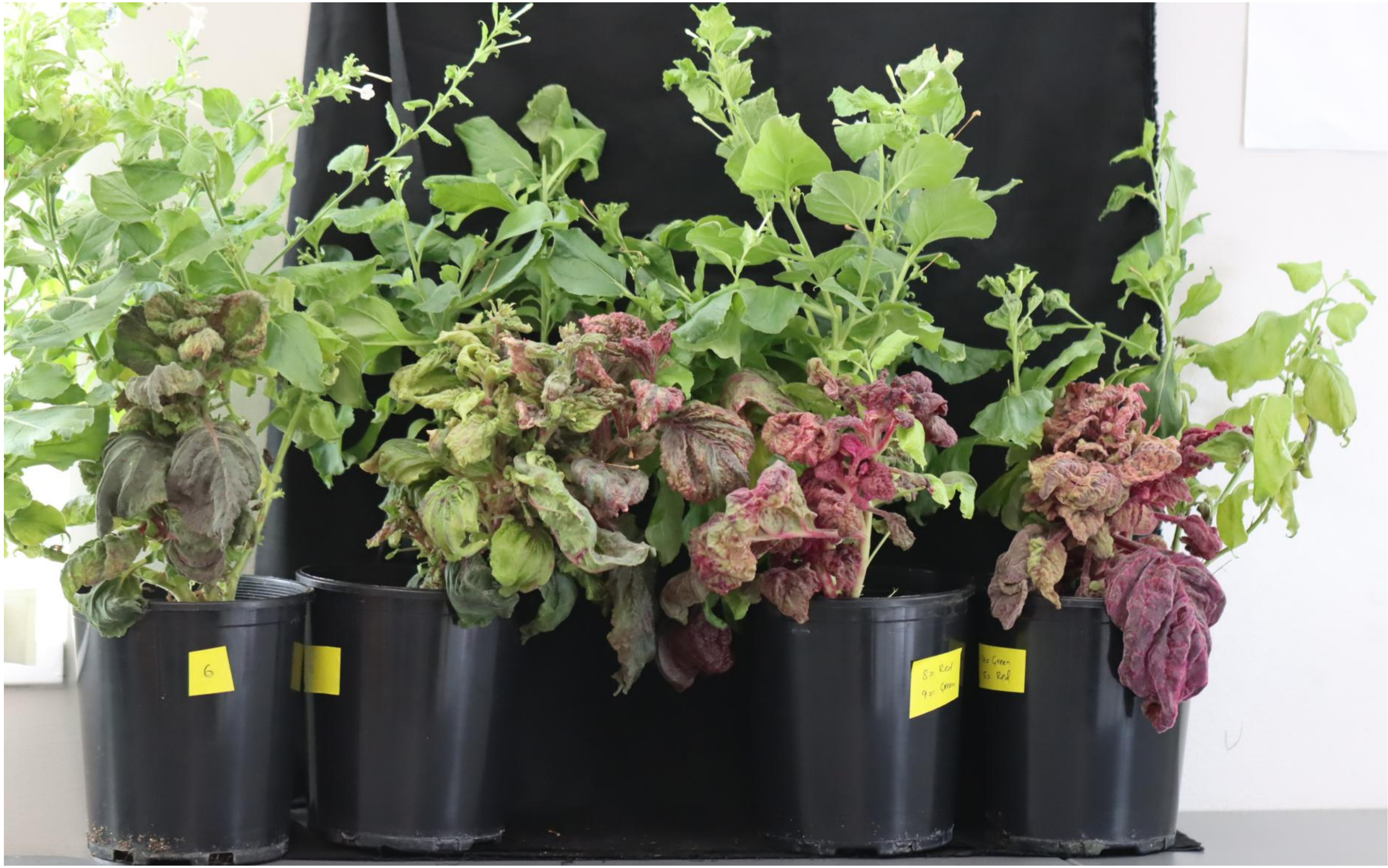

**Supplementary Fig. 3│** *In planta* transformation in *N. benthamiana* leading to regeneration of *RUBY* (betalain) expressing (red) shoots in the green mother plants. The representative image shows variable expression of *RUBY* in different T0 plants.

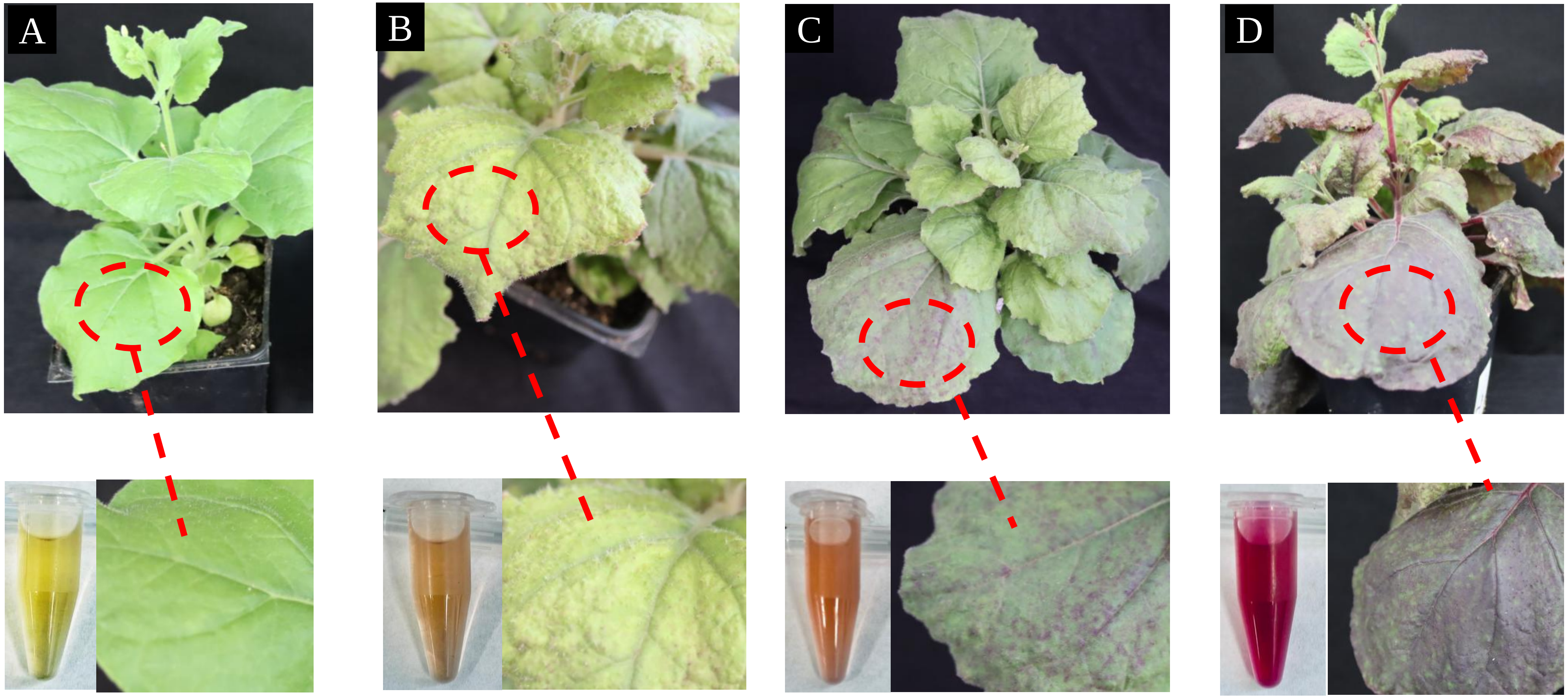


**Supplementary Fig. 4 | Transgenic shoots transmit the transgene to the progeny.** The plants in the progeny (T1) were analyzed for the transmission of transgenesis (A fragment of *RUBY* was PCR-amplified from the DNA extracted from these plants). However, the expression of RUBY was observed to vary, as shown in the figure above. Based on the pattern of *RUBY* expression in the progeny plants, they were scored as follows: panel **B 'Green'**, panel **C** as **'varigated'**, and panel **D** as **'Red'** with compared to the Wild Type, panel **A**. The tube show, juice samples extracted from 200mg of leaf tissue in 2 mL water. The intensity of betalain is shown for comparative purposes.

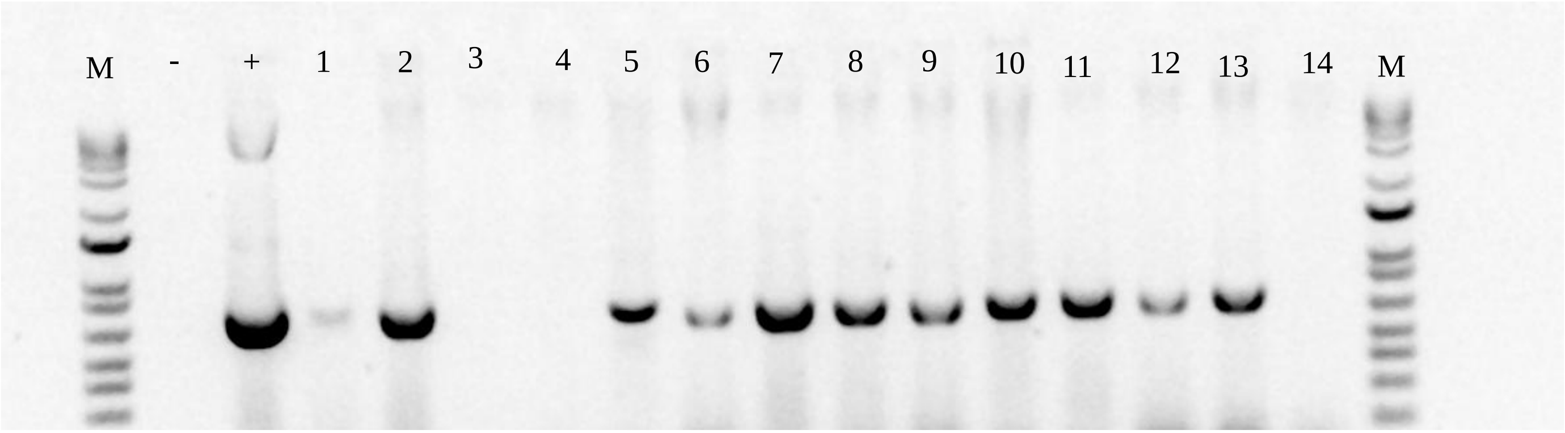


**Supplementary Fig. 5 |** PCR genotyping of representative *de novo* meristems, showing amplification of *ipt* to verify T-DNA insertion in tobacco (amplicon size 723bp). This verified the presence of T-DNA insertions in tobacco. (M= Marker; -= negative control and += positive control)

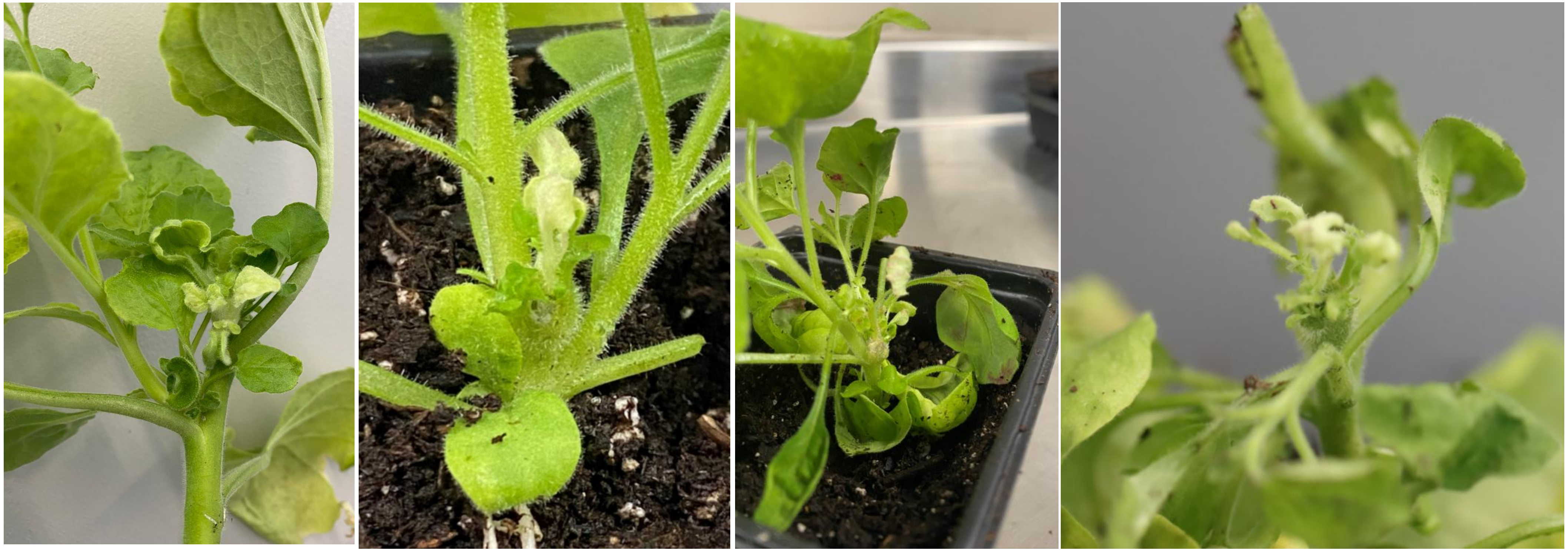

**Supplementary Fig. 6|** *In planta* transformation in tobacco with *WEipt1* construct induced regeneration of semi-albino shoots resulting from mono-allelic mutations in the (*PDS)* gene.

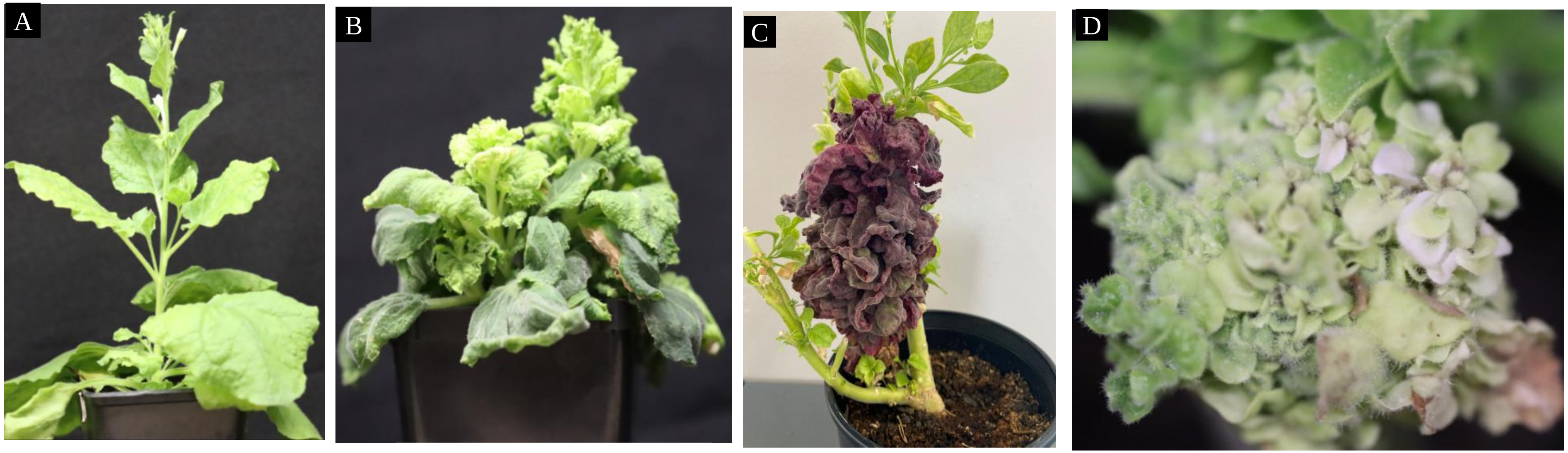


Normal stature (WT) | Abnormal stature | Abnormal stature/transgenesis | Abnormal stature/gene-editing

**Supplementary Fig. 7 |** Examples of *in planta* transformation in tobacco with *WEipt1* construct leading to the regeneration of *de novo* meristem with developmental defects in T0 generation. (**A**); WT. (**B-D**); abnormal stature.

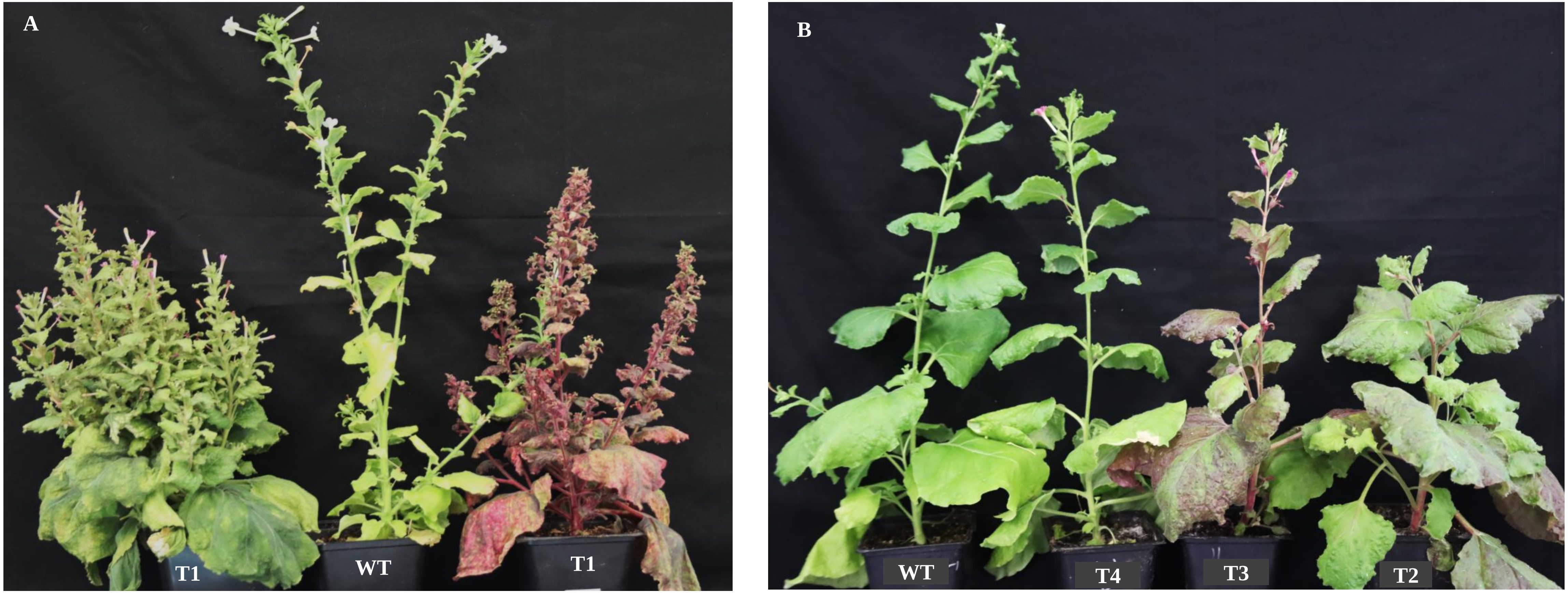


**Supplementary Fig. 8 |** Phenotypes of the progeny plants. The phenotype of T1 the plants was relatively normal except short stature, multiple branching, and flower setting  in comparison to WT **(A).** The phenotype of the plants was restored to normal in subsequent generations **(B).**

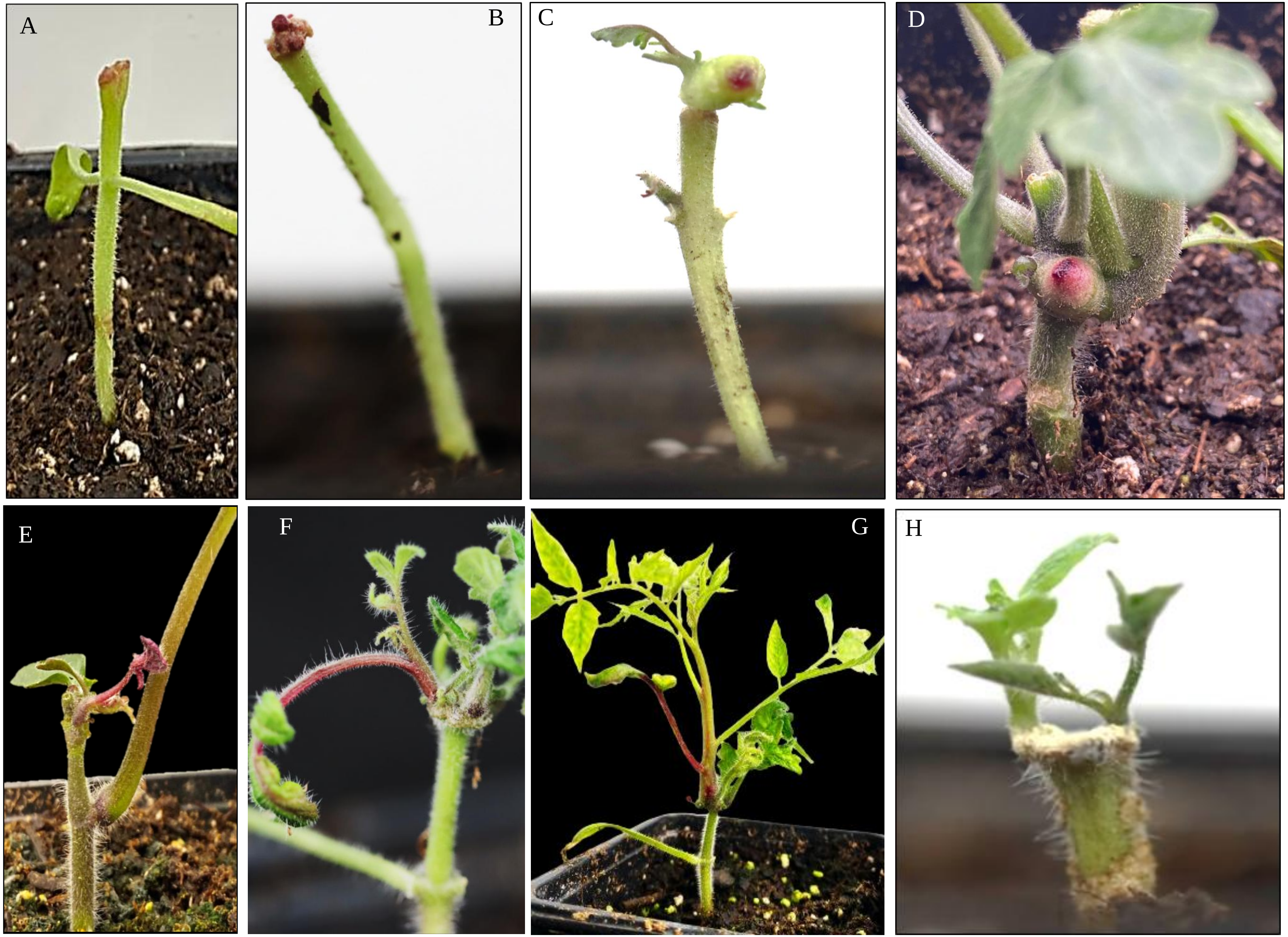


**Supplementary Fig. 9 |** *In planta* transformation of tomato with *WEipt* construct induced accelerated dedifferentiation and *de novo* meristem formation at the site of wounding. Transient expression of *RUBY*, five days post transfection **(A)** and ten days post-transfection **(B).** Regenerating callus with a non-transgenic (green) shoot and a transgenic callus **(C).** Transgenic callus that never regenerated **(D).** Regeneration of transgenic and non-transgenic shoots induced by *WEipt* (**E-H**).

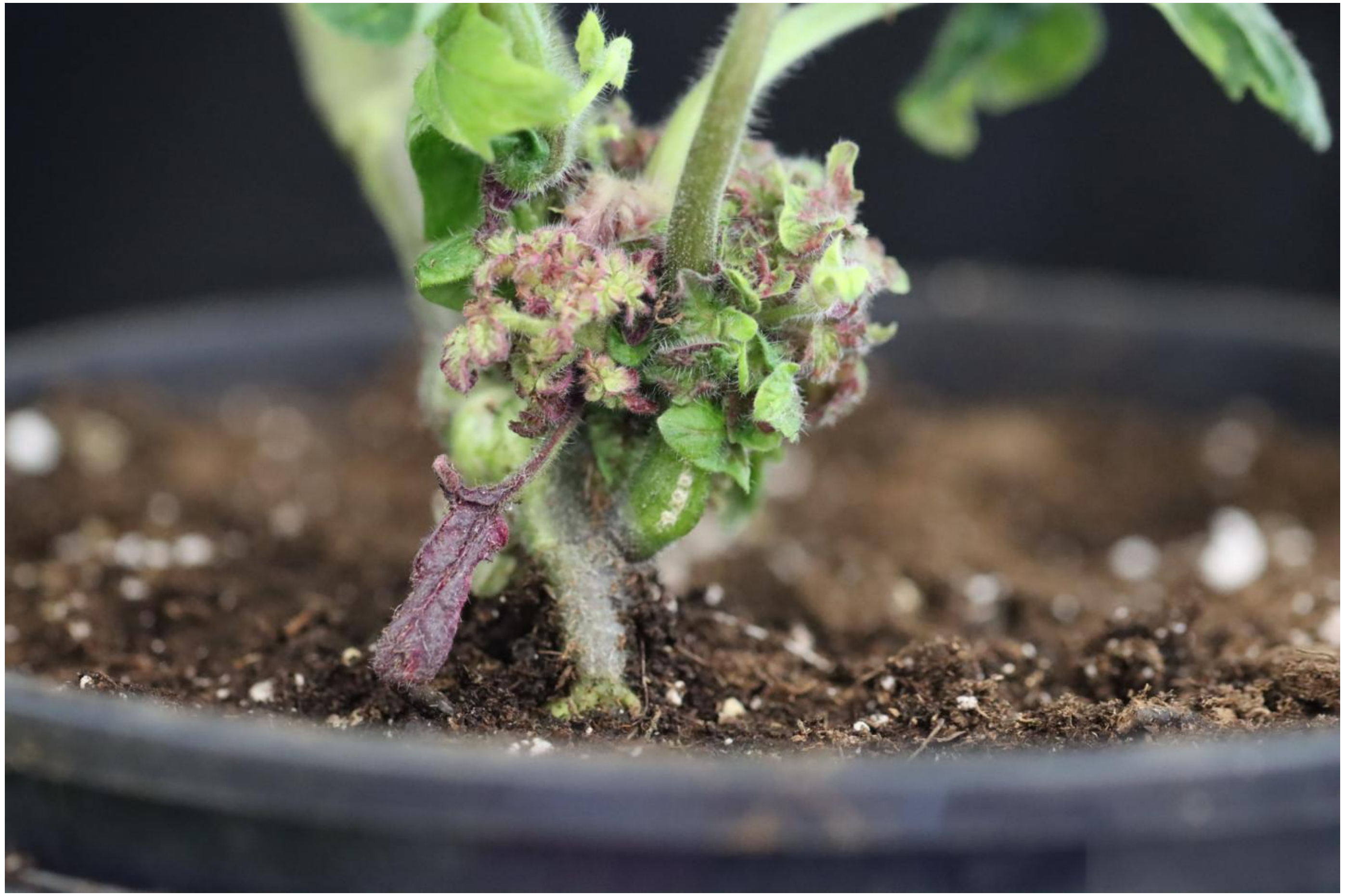

**Supplementary Fig. 10 |** Example of pleiotropic effects of *WEipt* observed in tomato in the T0 generation.

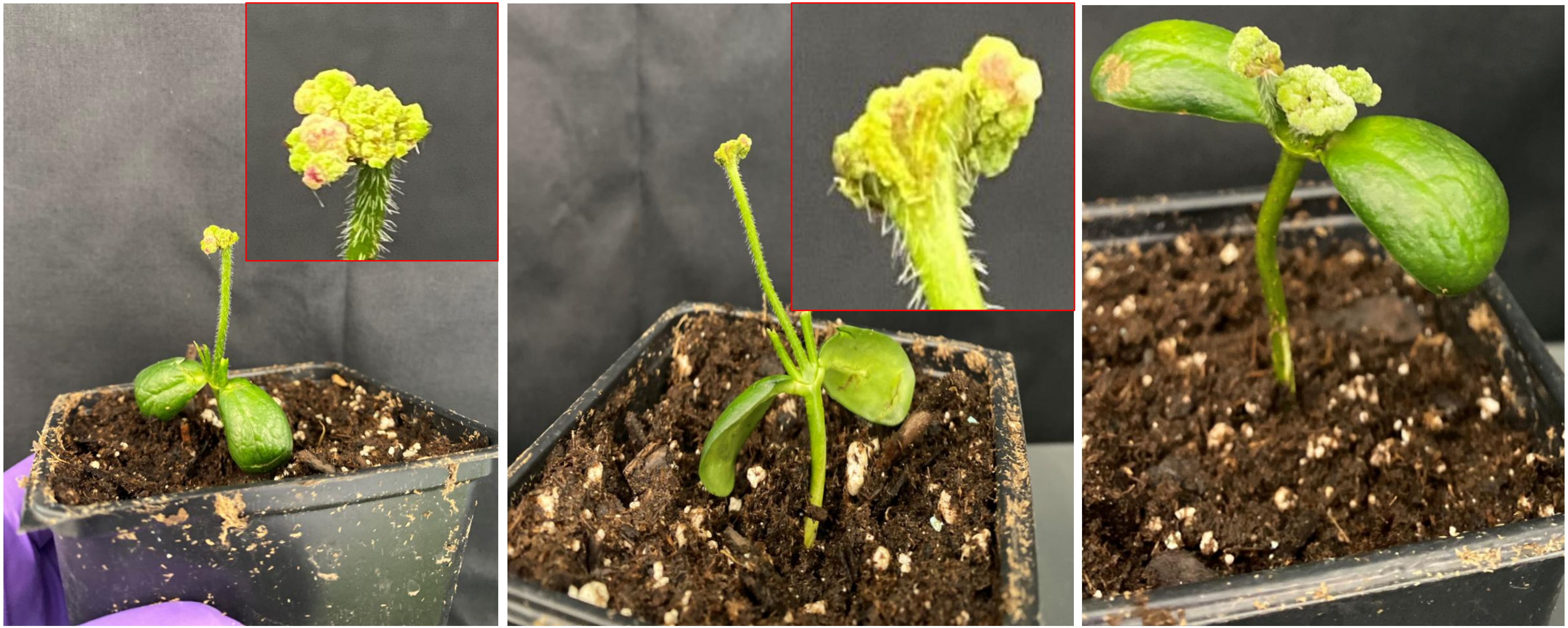

**Supplementary Fig. 11 |** *in planta* transformation of soybean with *WEipt2* led to rapid callus formation at the transfusion site without shoot regeneration. (Insets show images at larger scale).

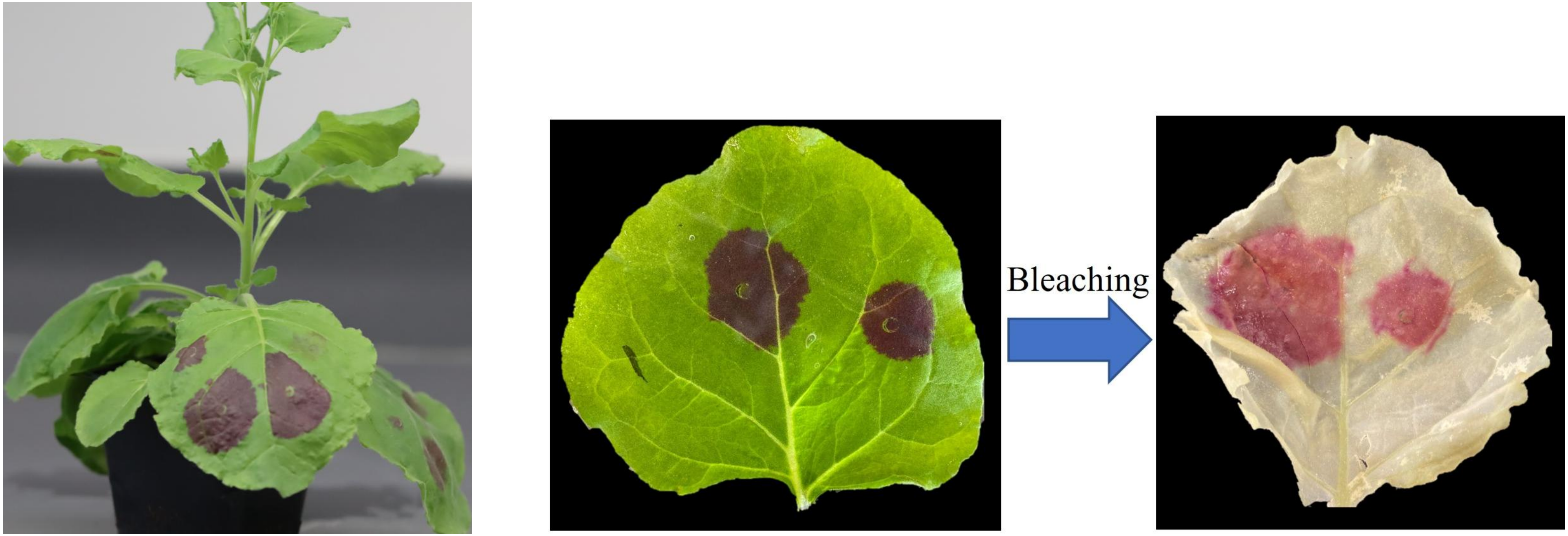


**Supplementary Fig 12 │** Visual confirmation of successful delivery of T-DNA carrying *RUBY* as a visible reporter gene, achieved through tobacco leaf agroinfiltration.
